# Light-dark dependent rhythmic changes in chloroplast and mitochondrial activity in *Chlamydomonas reinhardtii*

**DOI:** 10.1101/2025.04.28.650922

**Authors:** Gunjan Dhawan, Basuthkar Jagadeeshwar Rao

## Abstract

In photosynthetic organisms, inter-organellar coordination between mitochondria and chloroplast, particularly in synchronous cultures, has been widely appreciated but relatively less understood. Our study investigated the coordination in photosynthetic and mitochondrial activity during (12:12 h) light-dark cycle in *Chlamydomonas reinhardtii*. Live cell confocal imaging revealed light-dark-dependent mitochondrial morphology transitions from fragmented to intermediate to tubular forms by the end of 12 hr of light period, which reverses sharply through 6 and 12 hr of dark. Concurrently, chloroplast transitions from an intact cup (light) to a distorted and punctured structure (dark), which gets reversed in light phase. Spatial mapping showed tubular mitochondria positioned peripherally to the chloroplast cup in light, whereas fragmented and intermediate mitochondria were diffused around distorted chloroplast in dark, which again gets reversed in light. Functional analysis using 77K spectroscopy and photosynthetic protein levels (PsaA and D1) reflected that PSI/PSII fluorescence ratio remains stable in continuous light condition but increased exceptionally in continuous dark, which led to rhythmic oscillation in fluorescence ratio in light/dark-dependent manner in synchronous cultures. Mitochondrial activity, measured using Seahorse flux analyzer, showed basal oxygen consumption rate in continuous light and a marked reduction in continuous dark condition, resulting in rhythmic changes in light-dark cycle, indicating a coordinated rhythmicity in organellar function. Further, Target of rapamycin (TOR) kinase activity was essential to maintain inter-organellar coupled rhythmicity as evidenced by subdued rhythmicity following TOR kinase inhibition. The study, for the first time, argues for (12:12 h) light-dark cycle-mediated coupled rhythmicity between mitochondria and chloroplast in *C. reinhardtii*.

## Introduction

In photosynthetic organisms, daily alterations of light and dark regulate various physiological processes, including transcription, physiology, growth, and developmental processes (Harmer et al., 2009). These photosynthetic organisms try to adapt to the changing environmental conditions, and central to this adaptation are mitochondria and chloroplast, that play pivotal roles in energy metabolism.

Chloroplast functions as a primary site of carbon fixation by converting solar energy into biochemical energy through complex light-harvesting complexes and electron transport chain, which dynamically respond to the changing light conditions (Rochaix, 2011). Concurrently, mitochondrial metabolism, particularly the oxidative electron transport and phosphorylation processes are active in the light condition to maintain photosynthetic carbon assimilation (Raghavendra and Padmasree, 2003). Studies in plants have suggested mitochondria and chloroplast interaction at multiple levels, including the organic carbon exchange, regulation of ATP generation, carbon and nitrogen skeleton being exported from mitochondria and utilized for nitrogen assimilation in the chloroplast, and photorespiratory mechanisms (Yoshida and Noguchi, 2011).

Studies in photosynthetic algae have suggested that mitochondrial electron transport chain (ETC) plays a crucial role in retrograde regulation of the expression of genes related to photosynthesis (Matsuo and Obokata, 2006; Liu et al., 2024). The disruption of mitochondrial ETC function leads to changes in photosynthesis, particularly in electron transport and carbon metabolism (Cardol et al., 2003). On the other hand, mutation in photosynthetic pgrl1, which alters the cyclic electron flow, has shown to increase respiration rates in *C. reinhardtii* (Dang et al., 2014). Additionally, genetic mutation in the *Arabidopsis* multi-enzyme complex phosphoglycerate mutase/enolase/pyruvate kinase, involved in the exchange of metabolites between the two compartments, weakens the mitochondrial-chloroplast interactions (Zhang et al., 2020), while defects in *Arabidopsis* MICOS (mitochondrial contact site and cristae organizing system complex) components, disrupt the lipid trafficking between the two organelles, resulting in abnormal organellar morphology (Michaud et al., 2016; Li et al., 2019a). Overall, these studies suggest a case of organellar coupling between mitochondria and chloroplast in photosynthetic organisms. Therefore, we aimed to understand these dynamic changes in chloroplast and mitochondria in the light-dark cycle, using *C. reinhardtii* as a model system.

The unicellular algae *C. reinhardtii* serves as an excellent model for studying organelle dynamics during the diurnal cycle, as its cell division cycle is tightly synchronized with the diurnal cycle, such that the growth takes place in the light phase and division takes place in the dark (Zones et al., 2015). Along with this, *C. reinhardtii* shows the 12:12 hr light-dark cycle mediated rhythms in various biological processes such as chemotaxis, phototaxis, cell adhesion, starch content, cell division, nitrite uptake, sensitivity to UV irradiation (Matsuo and Ishiura, 2011). The cell cycle regulation of *C. reinhardtii* in synchrony has been extensively studied, but the impact of diurnal cycle on chloroplast and mitochondrial activity has received limited attention. Given the central role of these organelles in energy metabolism and cellular homeostasis, we investigated how the diurnal cycle influences the chloroplast and mitochondrial dynamics in *C. reinhardtii*.

The coordination of these organelle functions is crucial and is subjected to complex regulatory mechanisms. Among the most important and conserved regulatory pathways in eukaryotes is the TOR (target of rapamycin) kinase pathway. TOR kinase is a central node of nutrient sensing hub. It is a conserved serine/threonine protein kinase. In *C. reinhardtii,* TOR kinase is a central regulator of growth, development, metabolism, ER stress, autophagy, and lipid accumulation (Pérez-Pérez et al., 2017).

Recent studies in *Arabidopsis* have demonstrated the importance of TOR kinase in regulation of chloroplast activity, where TOR inactivation leads to downregulation of photosynthetic protein expression and biosynthetic pathways (Dong et al., 2015; Dobrenel et al., 2016b; Mallén-Ponce et al., 2022; D’Alessandro et al., 2024). Similarly, in mammalian systems, TOR kinase coordinates mitochondrial energy metabolism by stimulating mRNA translation and synthesis of nuclear-encoded mitochondria-related proteins, mitochondrial ribosomal proteins, and components of complex I and V (Morita et al., 2013). Collectively, these studies highlight the role of TOR kinase as a regulator of chloroplast and mitochondrial activity.

However, there is a void in literature regarding the role of TOR kinase in maintaining the balance between mitochondria and chloroplast in the diurnal cycle. The circadian clock may regulate the photosynthetic and respiratory functions, with TOR kinase being a mediator in response to external cues. Therefore, we tried to address the role of TOR kinase in the regulation of chloroplast and mitochondrial activities in a diurnal cycle.

In this study, we show that in synchronous culture, cells exhibited light-dark dependent morphological changes in mitochondria from fragmented to intermediate to tubular form by the end of the 12-hour of light period (onset of dark period), which reverses sharply through 6 and 12 hr of dark period. Concurrently, chloroplast shape changes from intact cup (12 hr light phase) into distorted and punctured structure in dark phase that in turn gets reversed through light phase. Spatial mapping revealed tubular mitochondria was apposed to the chloroplast cup peripherally in light conditions which changed to intermediate and fragmented mitochondria that were diffused throughout the distorted chloroplast cup in dark conditions which again gets reversed in light. Functional analyses using 77K spectroscopy and the photosynthetic protein levels (PsaA and D1 protein) reflected that F (PSI/PSII) ratio remains stable in the continuous light but increased exceptionally in the continuous dark condition. All these changes led to rhythmic oscillation in the fluorescence ratio in the light/dark-dependent manner in synchronous cultures. Concurrently, mitochondrial activity, measured using Sea horse flux analyzer, showed basal respiration rate in continuous light while the same in dark condition exhibited significantly lower oxygen consumption rate (OCR) resulting into rhythmic changes in 12:12 h light-dark cycle, suggesting a coordinated rhythmicity between chloroplast and mitochondrial function. Functional analyses using TOR kinase mutant show rhythmicity in chloroplast and mitochondrial functions similar to the wild-type cells, but the extent of rhythmicity is subdued. Taken together, our study highlights the coordinated rhythmicity in chloroplast and mitochondrial activities in *C. reinhardtii* where both the organelles are active in the light phase while their activity decreases in the dark phase, and the potential role of TOR kinase in maintaining the intensity of these organelle rhythms.

## Results

### Impact of diurnal cycle on the mitochondrial and chloroplast morphology in *C. reinhardtii*

To elucidate the impact of diurnal cycle on *C. reinhardtii* organelle morphology, we carried out live cell confocal imaging to monitor the changes in mitochondria and chloroplast. GFP-tagged mitochondrial strain (MDH4-GFP) was used to visualize different mitochondrial morphologies, and chloroplast was visualized using chlorophyll autofluorescence. Cells were cultured in three conditions: continuous light (LL), continuous dark (DD), and synchronous condition (LD), and imaged at different time points. We observed three distinct mitochondrial morphologies – tubular, intermediate, and fragmented, as reported previously (Rambold et al., 2011). In continuous light condition (LL), the mitochondrial morphology was predominantly tubular form, and the chloroplast cup remained intact (**Figure 1A** and **Video S1**). While in continuous dark condition, the mitochondria appeared as granular structures, and the chloroplast cup was distorted and punctured (**Figure 1A** and **Video S2**). In synchronous condition, mitochondria exhibited light-dark dependent morphology transition from fragmented to intermediate to tubular form by the end of the 12-hour of light period (**Figure 1B**, **Video S3A-S3C**), which reverses sharply upon dark incubation at 6 and 12 hr of dark (**Figure 1B**, **Video S3D and S3E**). These cells exhibiting different mitochondrial morphologies across all the time points were quantified and plotted as shown in **Figure 1C**, with the corresponding values provided in **Table S1A**. Moreover, the chloroplast cup showed punctate phenotype in the dark phase as opposed to the light phase (**Figure 1B**). All these results suggest that during synchrony, the cells containing tubular form of mitochondria increased as a function of light followed by a decrease of the same as the dark phase progresses.

**Figure 1:**
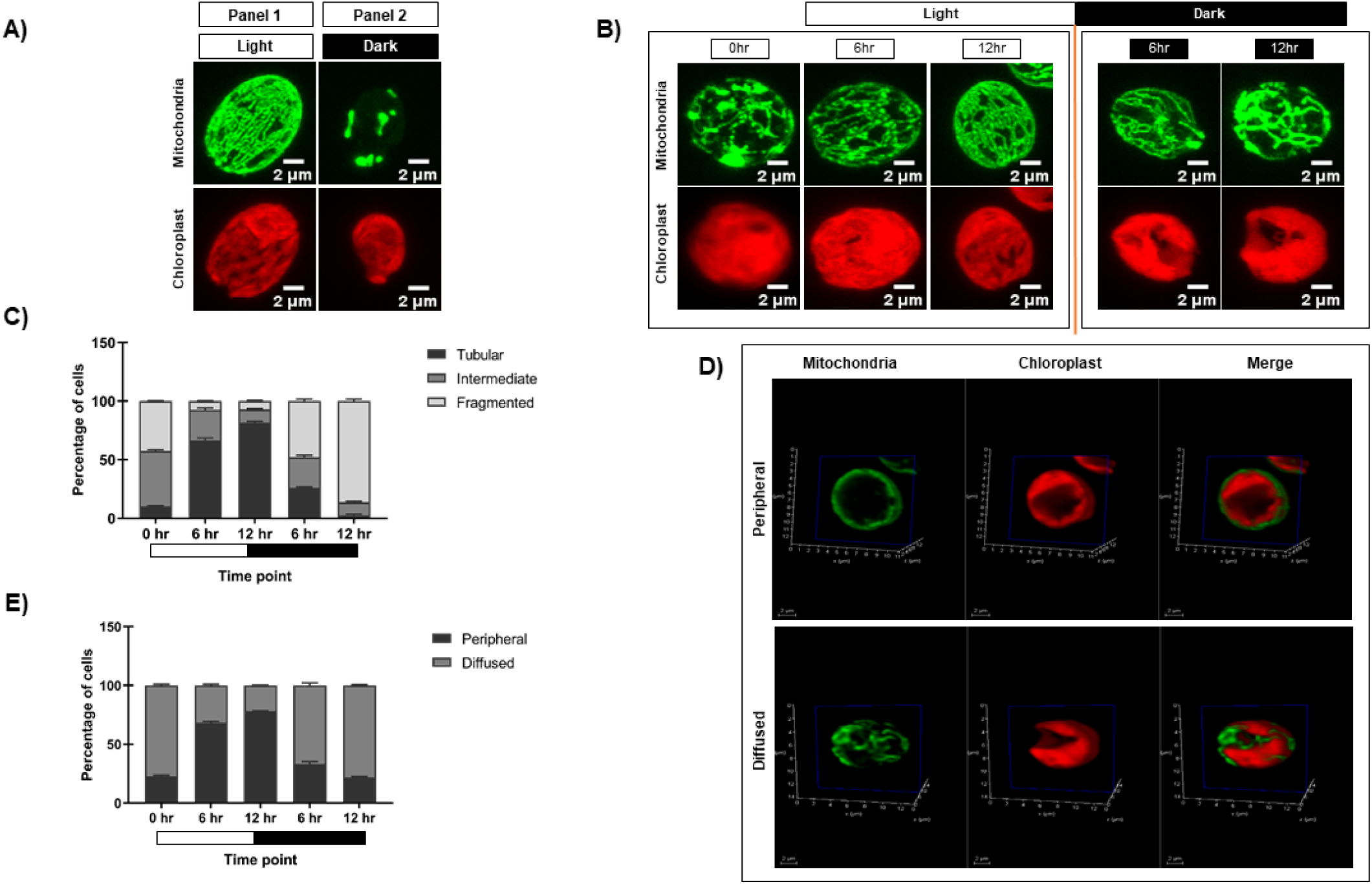
Morphological analysis of mitochondria and chloroplast in light and dark conditions. **(A)** Photoautotrophic *C. reinhardtii* mitochondrial GFP (MDH4-GFP) cells in continuous light (Panel 1) and continuous dark (Panel 2) were imaged using Leica TCS SP8 confocal laser scanning microscope and z-stack projected. MDH4-GFP cells were imaged for fluorescence at 488nm to visualize mitochondria. Chloroplast morphology was visualized by autofluorescence at 561nm. **(B)** *C. reinhardtii* cells following five days of 12:12 hour synchronization were imaged at 0, 6, and 12 hours of light (0 hr of dark) and 6 and 12 hours of dark period of the 24-hour cycle. **(C)** The percentage of cells having different mitochondrial morphologies at each time point was quantified using Image J software and plotted using GraphPad Prism 9.5.1 for MDH4-GFP cells. **(D)** Single plane of 3D rendering of mitochondria apposition with respect to chloroplast showing Peripheral (top panel) and Diffused (bottom panel) morphology. (**E)** Quantification of number of cells in peripheral and diffused arrangement of mitochondria apposition with respect to chloroplast for the MDH4-GFP cells.

In order to understand the relative position of mitochondria vis-à-vis chloroplast cup in the same samples, mitochondria and chloroplast were visualized on a single slice (**Figure 1D**). We observed 2 phenotypes: (1) mitochondria is along the periphery of the chloroplast (labeled as “peripheral”); (2) mitochondria is diffused throughout the cell (labeled as “diffused”). Upon quantification, it was observed that the peripheral phenotype of mitochondria vis-à-vis chloroplast cup gradually increased in the light phase of synchrony as opposed to the diffused phenotype (**Figure 1E** and **Table S1B**). Conversely, in the dark phase, the diffused phenotype of mitochondria was more prominent than the peripheral phenotype. 3D rendering was used to visualize the relative position of mitochondria vis-à-vis chloroplast cup as shown in **Videos S4** and **S5**. These results suggest that during synchrony, the peripheral association of mitochondria vis-à -vis chloroplast cup increases in the light phase followed by a decrease of the same in the dark phase, where the mitochondria diffuses in the entire cell.

Having established the clear organellar morphology dynamics in the light-dark cycle, we next analyzed whether these structural perturbations correlate with the changes in organellar functions by analyzing key photosynthetic and respiratory functional readouts.

### Analysis of PSI and PSII activity in the synchronous culture using 77K fluorescence spectroscopy

Photosynthetic activity was studied by measuring the relative abundance of photosystem I and II under LL, DD, and LD conditions. 77K fluorescence spectroscopy was performed, which is a measure of photosynthetic activity as reflected by PSI and PSII levels (Minagawa, 2011; Goldschmidt-Clermont and Bassi, 2015). At low temperature (77K), photosynthesis and non-photochemical quenching (NPQ) processes are inactive due to the lack of active proteins and thermal dissipation at this temperature. It allows fluorescence as the sole energy dissipation pathway, enabling more stable and distinct emission spectra for PSI and PSII (Lamb et al., 2015). Briefly, equal number of *C. reinhardtii* cells (3×10l cells/ml) were taken, and spectral readings were recorded at specified time points for each of the condition.

The samples grown in continuous light condition (LL) were collected at different time points (0, 6, 12, 18, and 24 hr), and 77K spectra were recorded (**Figure 2A**). Although these are representing the same samples, but we wanted to check if there is any time series effect. Two peaks were observed in the 77K spectra at around 685nm and 715nm, corresponding to PSII and PSI, respectively (Lamb et al., 2015). We observed relatively high PSI peak in relation to PSII at all the time points in continuous light condition (**Figure 2A**). The fluorescence ratio of PSI/PSII remained constant at all the time points (∼1.2-1.4) (**Figure 2A**, **2D** and **Table S2**). This ratio is similar to the previously reported F (PSI/PSII) ratio (Yadavalli et al., 2012). Similar to the continuous light condition, spectral readings were taken for the samples collected from the continuous dark condition (DD) at different time points (**Figure 2B**). These samples showed marked changes in the PSI/PSII fluorescence ratio, where PSII levels decreased in relation to PSI. PSI peak became broader and the resultant ratio of fluorescence from PSI to PSII increased significantly (∼1.7-2) (**Figure 2B, 2E** and **Table S2**). It suggests that the ratio of PSI/PSII is abnormally high in the continuous dark condition as opposed to the continuous light condition.

**Figure 2:**
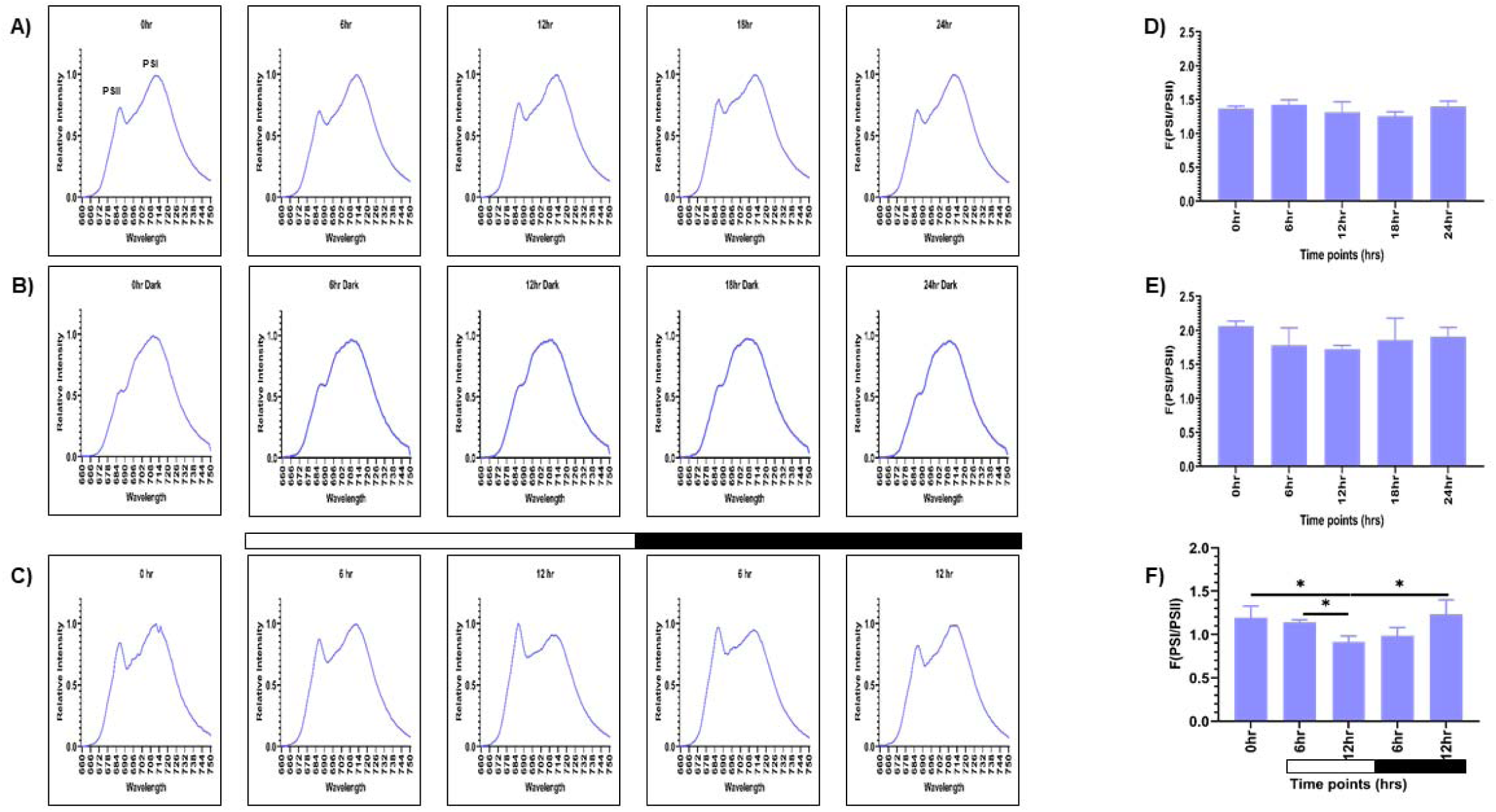
PSI and PSII spectral traces were recorded using 77K fluorescence spectroscopy. Average fluorescence emission spectral traces (77K spectra) were plotted for all three conditions: **(A)** continuous light (LL), **(B)** continuous dark (DD), and **(C)** 12:12 hour synchrony (LD) across all the time points. The ratio of F (PSI/PSII) for all three conditions **(D)** LL, **(E)** DD, and **(F)** LD at all the time points were plotted. The data represents an average of three independent biological repeats. Error bars represent Standard Deviation (SD). Statistical analyses were performed using 2way ANOVA.

Furthermore, to assess the photosynthetic activity in synchronous cultures, 77K spectra were recorded for the samples drawn at 0, 6, and 12 hr of light and 6 and 12 hr of dark (**Figure 2C**). There was a gradual decrease in the F (PSI/PSII) ratio in the light phase of synchrony, which increased concomitantly in the dark phase, approaching a value similar to the 0 hr sample (**Figure 2C, 2F** and **Table S2**). This experiment revealed that the relative activity of PSI and PSII are dynamically regulated in different light-dark phases of synchrony. Altogether, these findings suggest that the F (PSI/PSII) ratio remains stable in the continuous light condition but increases exceptionally in the continuous dark condition. In contrast, in synchronous cultures, there is rhythmic oscillation in the fluorescence ratio in the light/dark-dependent manner.

Further to corroborate these dynamic transitions observed in the 77K spectra, we checked for the protein level changes in PSI and PSII. The system was probed to analyze the levels of PsaA protein of PSI and PsbA (D1) protein of PSII, which are known markers of photosynthetic activity (Rochaix, 2002).

### Assessment of the photosynthetic proteins - PsaA and D1 upon synchrony

Western blot analysis was performed to observe the changes in levels of PsaA protein of PSI and PsbA (D1) protein of PSII, as a means to measure the photosynthetic activity. Briefly, total protein was extracted from samples drawn at specified time intervals for all three conditions, as described previously. We probed for D1 and PsaA proteins and used tubulin as the loading control. We observed constant levels of both D1 and PsaA proteins at all the time points in continuous light and continuous dark condition (**Figure 3A** and **3B**). However, the protein levels in the continuous dark condition were lower than that observed in the continuous light condition, indicating that there was an increase in the protein levels as a function of light and a decrease of the same in the dark condition. The protein levels were normalized using tubulin and expressed quantitatively, as shown in **Figure 3D** and **3E** (for D1 protein), and **Figure 3G** and **3H** (for PsaA protein) for LL and DD condition, respectively with the corresponding values provided in **Figure 3J** and **3K**, respectively.

**Figure 3:**
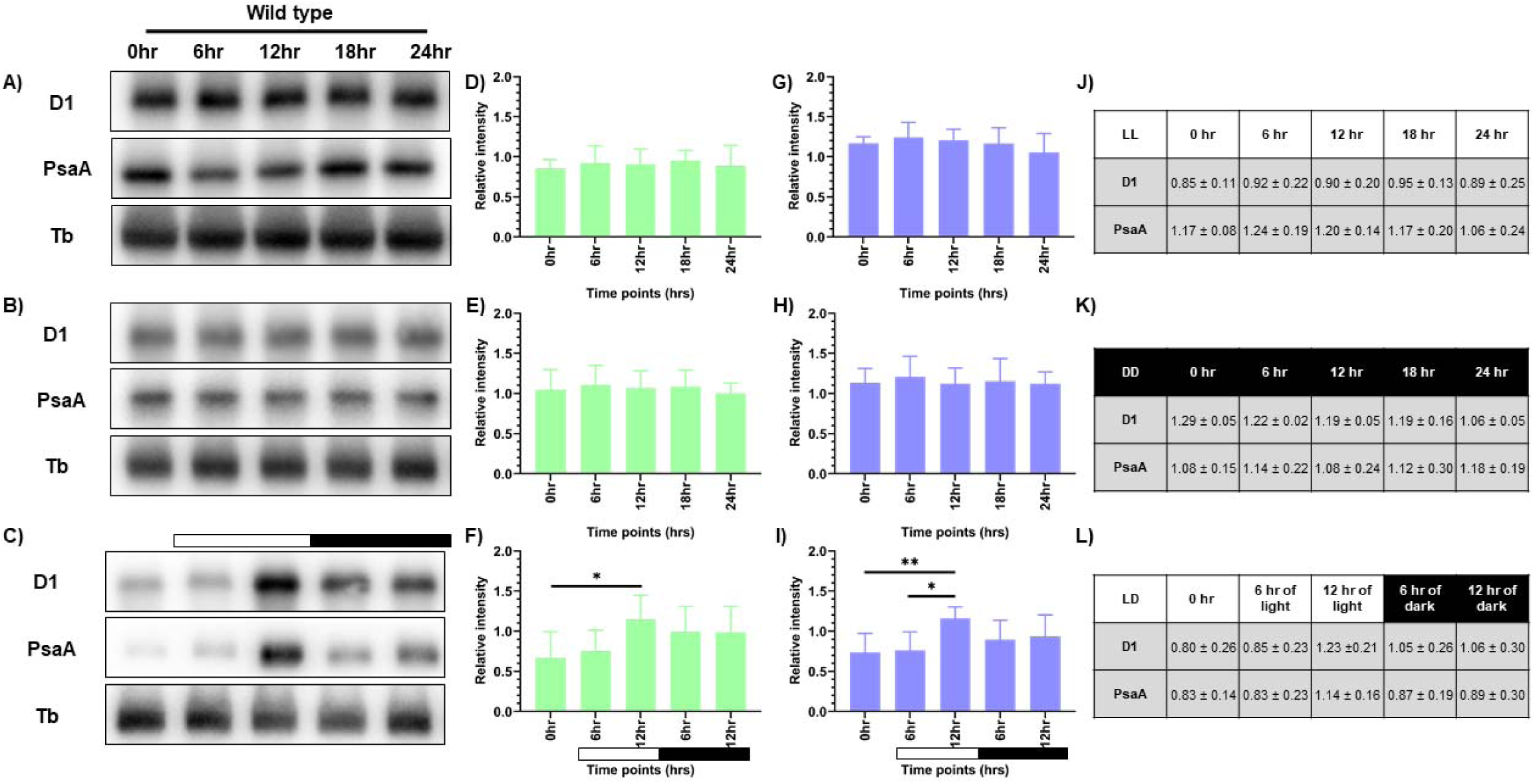
Expression levels of photosynthetic proteins in the wild-type cells. Western blot analysis of PsaA and PsbA/D1 proteins in wild-type cells at different time points of the 24-hour cycle were shown for **(A)** continuous light, **(B)** continuous dark, and **(C)** 12:12 hour synchrony condition. The relative protein levels were quantified by measuring the D1 and PsaA band intensity with respect to tubulin. Panel **D-F** shows the expression of D1 protein, and panel **G-I** shows the levels of PsaA protein in LL, DD, and LD conditions, respectively, with the corresponding values shown in panel **J-L**. The data represents an average of three independent biological repeats. Error bars represent Standard Deviation (SD). Statistical analyses were performed using 2way ANOVA.

In the synchronous culture, abundance of both the proteins gradually increased during the light phase, reaching a maximum at 12 hr of light (**Figure 3C**). Meanwhile, in the dark phase, levels of both the proteins decreased (**Figure 3C**). The protein levels were normalized and quantitatively expressed in **Figure 3F** (for D1 protein) and **3I** (for PsaA protein), with the corresponding values provided in **Figure 3L**. The increase in the relative levels of D1 and PsaA proteins, as a function of light and decrease of the same in the dark, reflects the light-dark dependent photosynthetic changes in the *C. reinhardtii* cells. These transient changes in the levels of photosynthetic proteins coincide with the rhythmic changes in the relative PSI to PSII fluorescence, observed in the 77K spectral data, in the light-dark cycle (**Figure 2C** and **2F**).

These dynamic changes in the 77K spectra and the photosynthetic protein levels reflect rhythmicity in the photosynthetic activity as a function of light-dark cycle, thus prompting us to examine whether similar transitions are evident in case of mitochondrial functions.

### Changes in respiratory oxygen consumption upon synchrony

To evaluate the mitochondrial activity, we measured mitochondrial respiration rate. Seahorse XFp Flux analyzer was used to provide the intracellular readout of oxygen consumption rate [OCR (pmol/min per 1×10^6^ cells)] at 24lC in the dark as a function of time. We measured the coupled respiration, which results in generation of ATP (reflecting basal respiration rate) and uncoupled respiration (using 1 µM CCCP), which is associated with the dissipation of excess proton gradient as heat (reflecting maximal respiration rate). Briefly, samples from all three conditions – LL, DD, and LD were drawn, as described previously. These samples were incubated in dark for 20 minutes prior to the first reading, to nullify the photosynthetic oxygen changes. The average of OCR values at the first three time points in the graph depict basal respiration rate (**Figure 4A**). Then, the mitochondrial uncoupler CCCP is injected in order to get the maximal respiration rate. However, unlike mammalian cells, in *C. reinhardtii*, the surge in maximal OCR is transient, and it drops down after attaining a maximal value.

**Figure 4:**
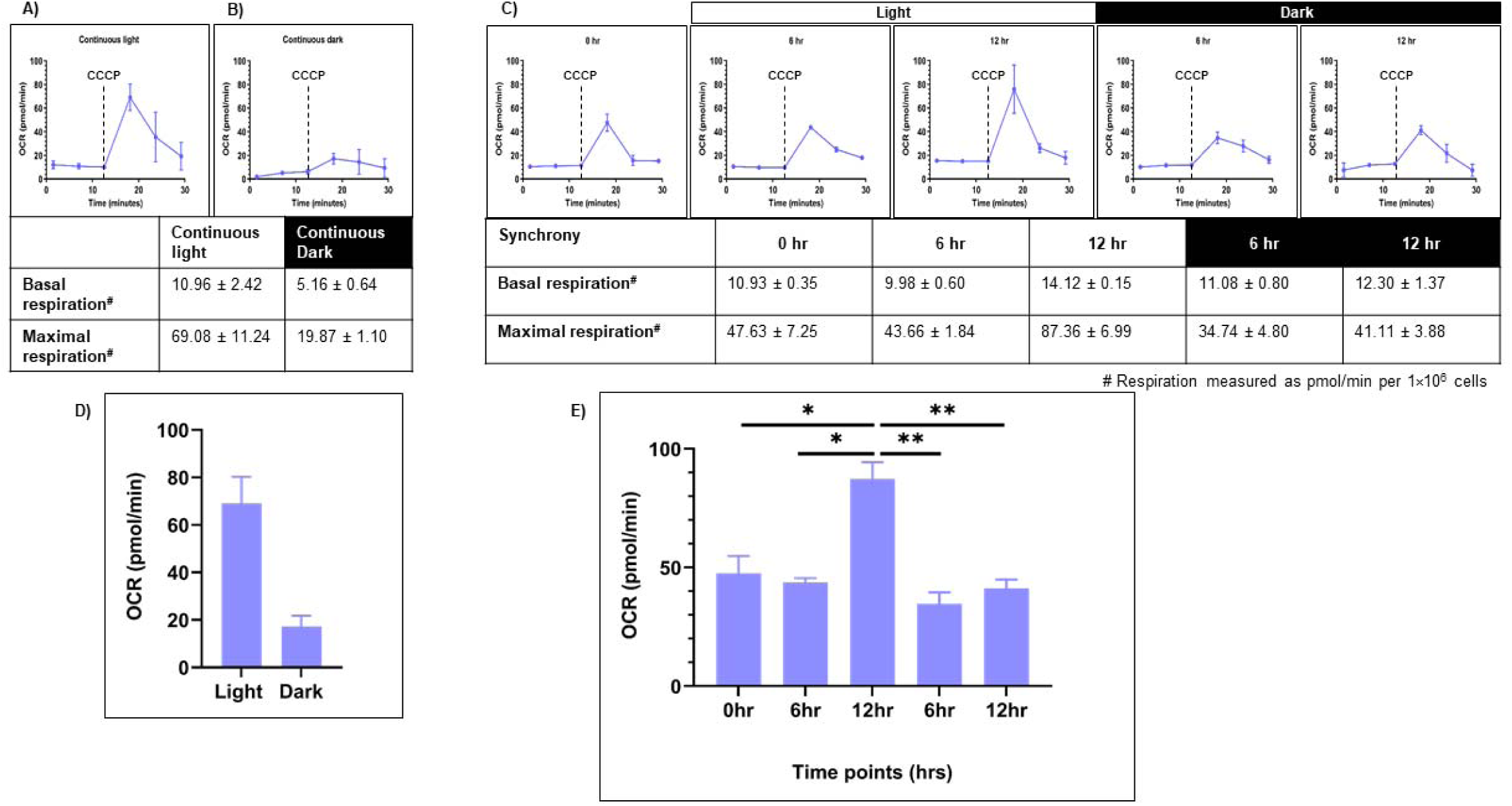
Mitochondrial respiration rate was measured for the wild-type cells. Seahorse Flux analyzer assay was performed for the oxygen consumption rate (OCR) measurements for all three conditions: **(A)** continuous light (LL), **(B)** continuous dark (DD), and **(C)** 12:12 hr synchrony (LD). **(D)** The maximal respiration after the addition of CCCP was plotted for wild-type cells in continuous light and continuous dark condition and **(E)** synchronous culture. The data represents an average of three independent biological repeats. Error bars represent Standard Deviation (SD). Statistical analyses were performed using 2way ANOVA.

Cells grown under continuous light condition (LL) were subjected to OCR measurements (**Figure 4A**). These samples showed the basal respiration rate of 10.96 ± 2.42 (all values represent mean ± SD, n=3). Upon CCCP addition, the value increased to 69.08 ± 11.24 (**Figure 4A** and **4D**). In contrast, the cells drawn from the continuous dark condition exhibited significantly lower basal OCR (5.16 ± 0.64) compared to the continuous light condition (**Figure 4B**). Moreover, following CCCP addition, maximal respiration rate is 19.87 ± 1.10, which is low in comparison to the continuous light condition (**Figure 4B** and **4D**). This data indicates that the mitochondrial activity is attenuated in the dark-grown cultures as opposed to the light-grown cultures.

Further, in the synchronous culture, the basal respiration rate remained constant across all the time points of synchrony (**Figure 4C**). These OCR values were comparable to that observed in the continuous light condition (**Figure 4A**). However, following the addition of CCCP, there was a significant increase in OCR levels in the light phase, reaching a maximum value at 12 hr of light, followed by a concomitant decline of the same during the dark phase of synchrony (**Figure 4C** and **4E**). These transient changes in the oxygen consumption rate suggest the rhythmicity in mitochondrial activity, which aligns with the rhythmic changes observed in photosynthetic activity in the light-dark phases of synchrony.

Overall, our study elucidates the rhythmic pattern in chloroplast and mitochondrial activity in the light-dark cycle. Further, we explored the potential role of TOR kinase in regulating these organelle rhythms. Given the conserved role of TOR kinase in cellular growth, development and metabolism (Morita et al., 2013; Dobrenel et al., 2016a), we aimed to understand whether TOR kinase contributes to the organellar rhythmicity in the light-dark cycle. For that, cells were probed with TOR kinase inhibitor to observe the morphological changes in these organelles.

### Impact of TOR kinase inhibition on mitochondrial and chloroplast morphologies in synchronized cultures

To gain insight into the effect of TOR kinase inhibition, we employed a classical TOR kinase domain inhibitor AZD8055 (Chresta et al., 2010; Dong et al., 2015), which directly targets the catalytic domain of TOR kinase by competing with ATP in its binding pocket, thus inhibiting the kinase activity. TOR kinase inhibition is assayed by monitoring the changes in phosphorylation levels of its downstream substrate CrS6K (Pérez-Pérez et al., 2017; Upadhyaya et al., 2020). We observed the reduction in CrS6K phosphorylation levels in the AZD-treated cells compared to the control cells (**Figure S1**), which confirmed the efficiency of TOR kinase inhibitor in our current experimental conditions.

After validating the inhibition protocol, chloroplast and mitochondrial morphologies were examined in the TOR kinase-inhibited cells upon synchrony. Live cell confocal imaging was performed, as described previously, to visualize the chloroplast and mitochondria. The predominant mitochondrial and chloroplast morphologies obtained at different time points of synchrony in the TOR kinase-inhibited cells are illustrated in **Figure 5A** and **Video S6A-S6E**. In synchronous condition, the tubular form of mitochondria increased on exposure to light and then decreased in the dark in the TOR kinase-inhibited cells (**Figure 5A**), similar to the control cells (**Figure 1B**). However, while control cells showed the maximum of tubular morphology at 12 hours of light, the inhibited cells achieved the peak at only 6 hours of light before starting to decrease (**Figure 5A** and **5B**). Whereas, during the dark phase, both cell types predominantly exhibited fragmented and intermediate forms of mitochondria. The TOR kinase-inhibited cells exhibiting different mitochondrial morphologies across all the time points were quantified and plotted as shown in **Figure 5B**, with the corresponding values given in **Table S3A**. Moreover, unlike control cells (**Figure 1B**), which exhibited an intact chloroplast cup during the light phase and distortion of the same in the dark phase, in the TOR kinase-inhibited cells, chloroplast distortions were persistent throughout the light-dark cycle (**Figure 5A** and **Video S6A-S6E**). Altogether, these results suggest that upon TOR kinase inhibition, the tubular form of mitochondria increases as a function of light and drops in the dark phase, similar to the control cells. However, there is subdued rhythmicity in case of TOR kinase-inhibited cells compared to the control cells.

**Figure 5:**
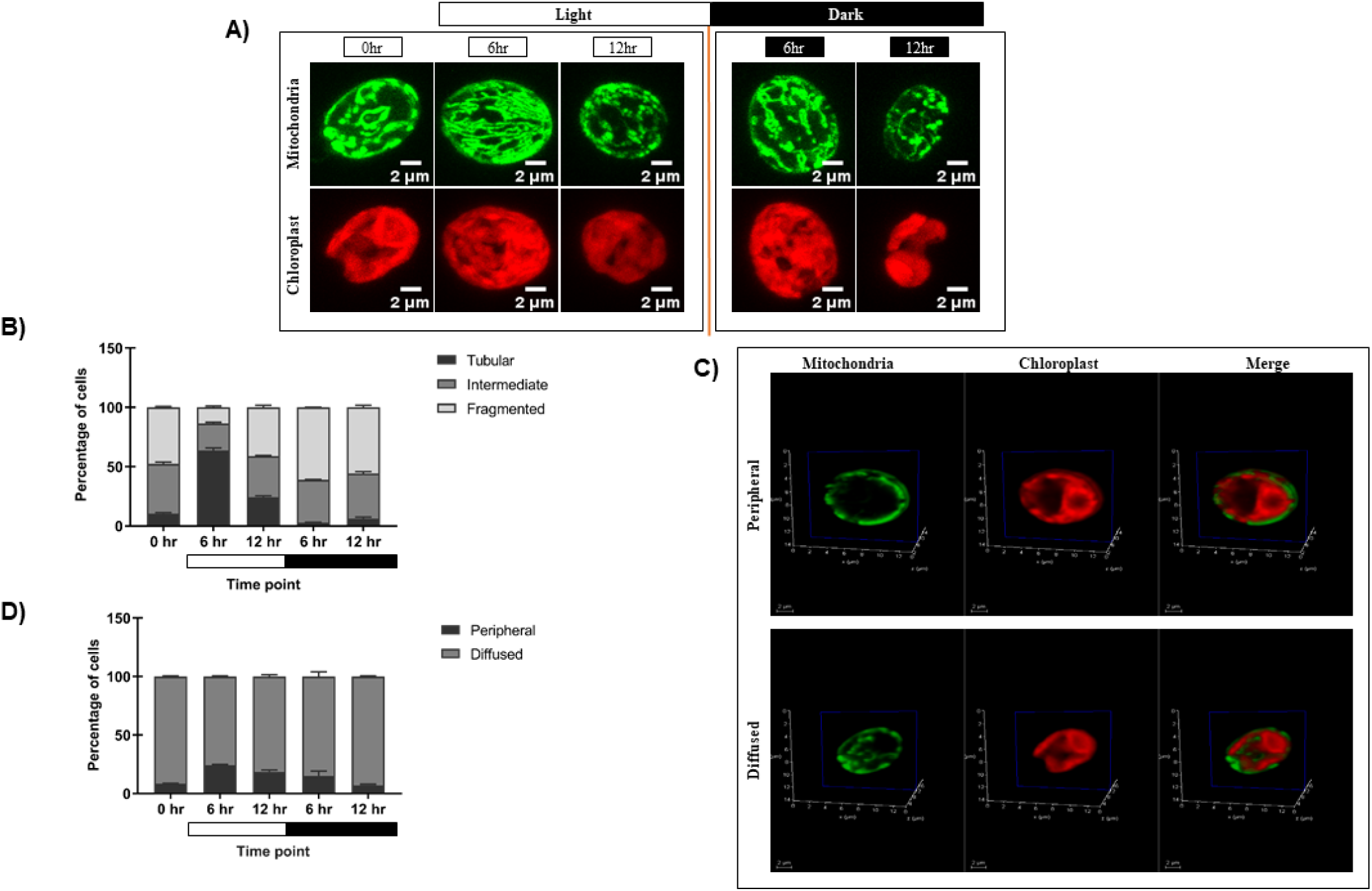
Effect of TOR kinase inhibition by AZD8055 treatment on the mitochondrial and chloroplast morphology in the light-dark cycle. **(A)** *C. reinhardtii* mitochondrial GFP (MDH4-GFP) cells following 12:12 hr synchronization were treated with AZD8055 (1µM) for achieving the TOR kinase inhibition and imaged at 0, 6, and 12 hours of light (0 hr of dark) and 6 hr and 12 hr of dark period for 24 hours. Mitochondria (green) and chlor oplast (red). **(B)** Quantification of number of AZD8055 treated cells having different mitochondrial morphologies at each time point as obtained in panel a. **(C)** Images in panel a were analyzed at a single plane of 3D rendering of mitochondria apposition with respect to chloroplast showing Peripheral (top panel) and Diffused (bottom panel) morphology. **(D)** Quantification of the percentage of cells containing peripheral and diffused morphology upon TOR kinase inhibition.

Furthermore, the relative position of mitochondria vis-à-vis chloroplast in the TOR kinase-inhibited cells was visualized in the same samples by 3D rendering (**Figure 5C** and **Video S7** and **S8**). Unlike control cells, which showed a clear transition in the peripheral to diffused phenotype in the light-dark phases of synchrony (**Figure 1E**), the majority of TOR kinase-inhibited cells exhibited diffused phenotype at all the time points of synchrony (**Figure 5D** and **Table S3B**), thus suggesting that the transition is compromised in case of TOR kinase inhibited cells.

Then, we assessed the functional implications of these morphological changes observed in the TOR kinase-inhibited cells and compared the same with the control cells. To avoid time-dependent perturbations associated with the incubation step of TOR kinase inhibitor, we used a TOR kinase insertional mutant (LMJ.RY0402.203031) for subsequent analyses of photosynthetic and respiratory functions (Li et al., 2019b).

### Analysis of the PSI and PSII activity in the synchronous culture of TOR kinase mutant cells

We assessed the changes in photosynthetic activity in the TOR kinase mutant using 77K spectroscopy. Spectral readings were recorded at specified time points for each of the condition (LL, DD, LD), as described previously for the wild-type cells. In continuous light condition, the TOR kinase mutant recapitulated the wild-type PSI/PSII stoichiometry (**Figure 2A**), displaying high PSI level in relation to PSII, at all the time points (**Figure 6A, 6D** and **Table S4**). Moreover, in the continuous dark condition, mutant cells exhibited a broadened PSI peak, and the resulting F (PSI/PSII) ratio was around 1.7-2.1 (**Figure 6B, 6E** and **Table S4**), which was comparable to the wild-type cells (**Figure 2B**). Collectively, these results suggest that the fluorescence ratio in TOR kinase mutant remains similar to that of wild-type cells in both LL and DD condition.

**Figure 6:**
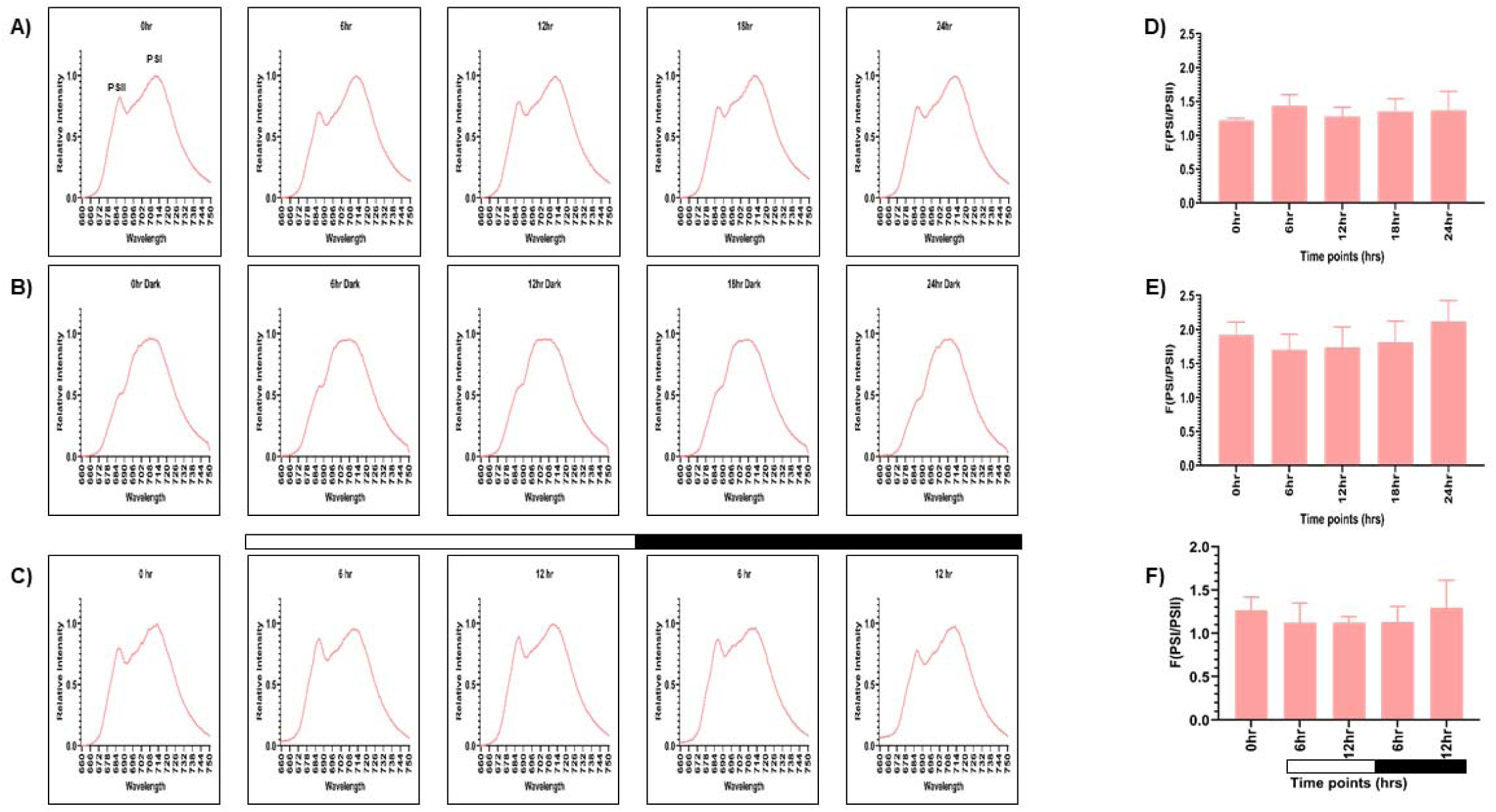
Fluorescence emission spectra using 77K spectroscopy were recorded for the TOR kinase mutant. Average fluorescence emission spectral traces (77K spectra) were plotted for all three conditions: **(A)** continuous light (LL), **(B)** continuous dark (DD), and **(C)** 12:12 hour synchrony (LD) across all the time points for the TOR kinase mutant. The ratio of F(PSI/PSII) for all three conditions **(D)** LL, **(E)** DD, and **(F)** LD for the TOR kinase mutant at all the time points were plotted. The data represents an average of three independent biological repeats.

Furthermore, in the synchronous condition, similar to our earlier observations in the wild-type cells (**Figure 2C**), the F (PSI/PSII) ratio in the mutant cells exhibited rhythmic changes, where the ratio decreases in the light phase and increases in the dark phase (**Figure 6C, 6F** and **Table S4**). However, this rhythmic change is less pronounced in the TOR kinase mutant compared to the wild-type cells. To validate these changes in the photosynthetic activity, we analyzed the levels of photosynthetic proteins in the TOR kinase mutant.

### Analysis of the photosynthetic proteins - PsaA and D1 in the TOR kinase mutant upon synchrony

Similar to the wild-type condition, western blot analysis was performed to probe for D1 and PsaA protein in the TOR kinase mutant. Consistent with the wild-type cells (**Figure 3A** and **3B**), TOR kinase mutant showed higher levels of both D1 and PsaA protein in continuous light condition compared to the continuous dark condition (**Figure 7A** and **7B**) and the protein levels remained constant over 24 hours period. The protein levels were normalized for both – LL and DD conditions and expressed quantitatively in **Figure 7D** and **7E** (for D1), **7G** and **7H** (for PsaA), respectively, and the corresponding values are given in **Figure 7J** and **7K**, respectively. These findings suggest that protein content in TOR kinase mutant is comparable to the wild-type cells in both continuous light and dark condition.

**Figure 7:**
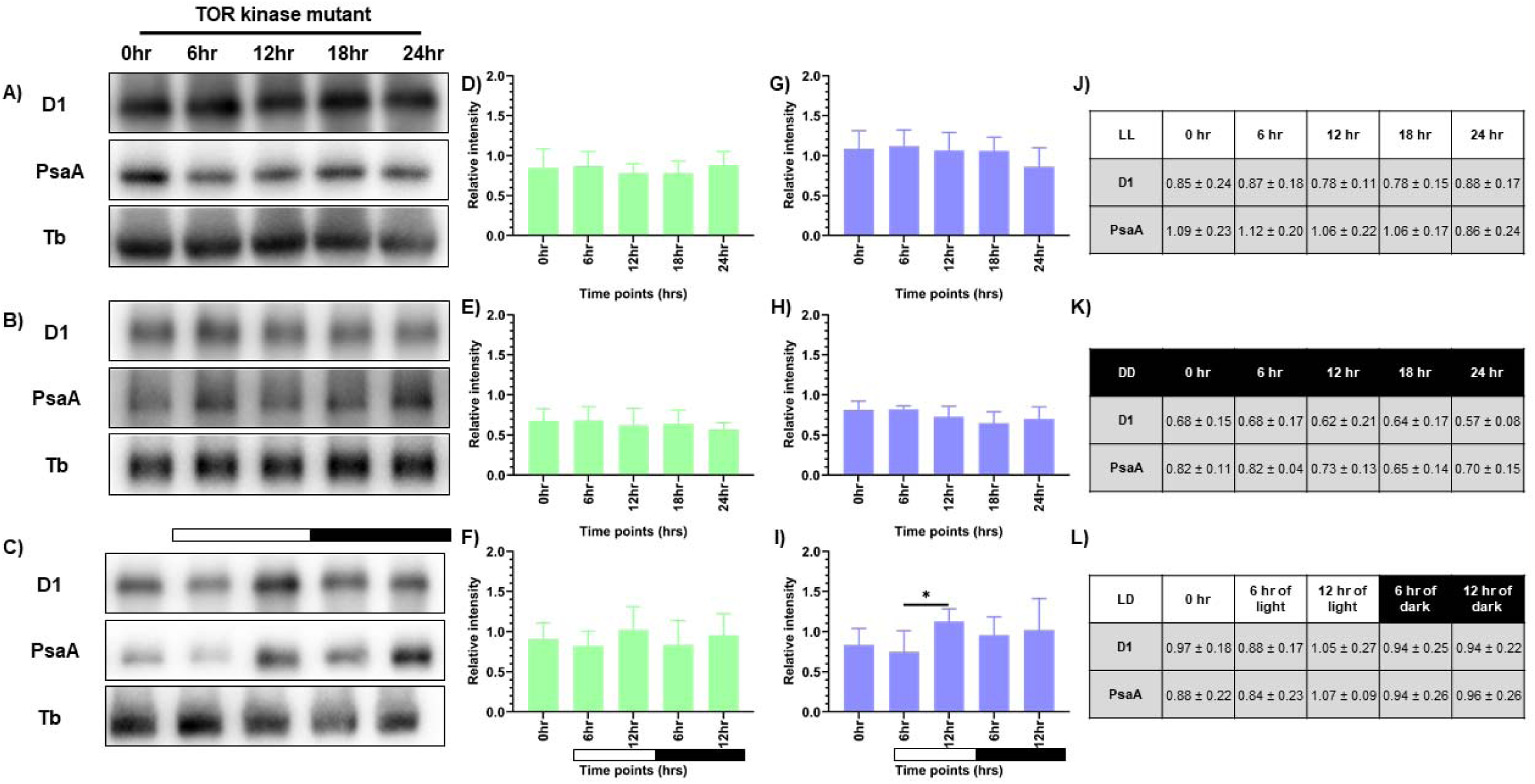
Expression levels of photosynthetic proteins in the TOR kinase mutant. Western blot analysis of PsaA and PsbA/D1 proteins in the TOR kinase mutant at different time points of the 24-hour cycle were shown for **(A)** continuous light, **(B)** continuous dark, and **(C)** 12:12 hour synchrony condition. The relative protein levels were quantified by measuring the D1 and PsaA band intensity with respect to tubulin in the TOR kinase mutant. Panel **D-F** shows the expression of D1 protein, and panel **G-I** shows the levels of PsaA protein in LL, DD, and LD conditions, respectively, in the TOR kinase mutant, with the corresponding values shown in panel **J-L**. The data represents an average of three independent biological repeats. Error bars represent Standard Deviation (SD). Statistical analyses were performed using 2way ANOVA.

In synchronous condition, similar to the wild-type cells (**Figure 3C**), we observed a rhythmic change in protein levels in the TOR kinase mutant (**Figure 7C**), where the protein levels increased as a function of light and decreased in the dark. However, unlike wild-type cells, the rhythmicity in the TOR kinase mutant is less pronounced. The quantitative analyses of D1 and PsaA proteins are depicted in **Figure 7F** and **7I**, respectively, with the corresponding values shown in **Figure 7L**. These results indicate subdued protein-level rhythmicity in the TOR kinase mutant compared to the wild-type cells. This data is consistent with the attenuated 77K spectral rhythmicity, indicating overall subdued photosynthetic activity in the TOR kinase mutant compared to the wild-type cells. Given this photosynthetic rhythmicity, we next assessed whether mitochondrial functions were similarly affected in the diurnal cycle in TOR kinase mutant.

### Changes in respiratory oxygen consumption in the TOR kinase mutant upon synchrony

The OCR measurements were performed to analyze the mitochondrial activity in TOR kinase mutant, as described previously for the wild-type cells. In the continuous light condition (LL), similar to the wild-type cells (**Figure 4A**), in TOR kinase mutant we recorded a basal level of respiration (15.61 ± 3.86), which markedly increased to 78.21 ± 5.44 upon the addition of CCCP (all values represent mean ± SD, n=3) (**Figure 8A** and **8D**). In contrast, in the continuous dark condition, we observed low basal respiration rate (4.08 ± 0.64) and maximal respiration rate (39.93 ± 13.08), compared to the continuous light condition (**Figure 8B** and **8D**). It suggests a trend similar to the wild-type cells (**Figure 4D**), where the OCR values were higher in the continuous light condition compared to continuous dark condition.

**Figure 8:**
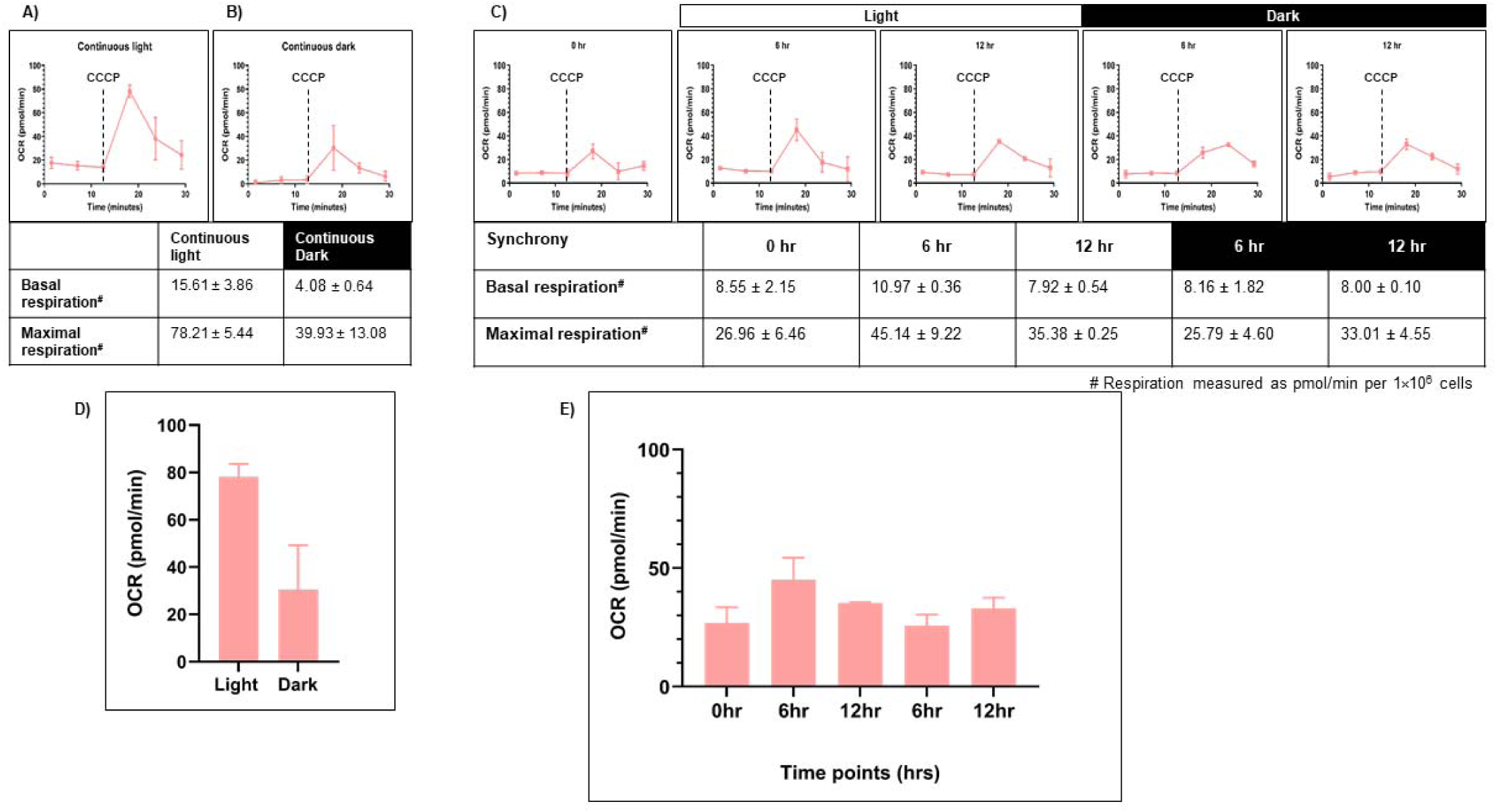
Mitochondrial respiration rate was measured for TOR kinase mutant. Seahorse Flux analyzer assay was performed for the oxygen consumption rate (OCR) measurements for all three conditions: **(A)** continuous light (LL), **(B)** continuous dark (DD), and **(C)** 12:12 hr synchrony (LD) for the TOR kinase mutant. **(D)** The maximal respiration after the addition of CCCP was plotted for the TOR kinase mutant in continuous light and continuous dark condition and **(E)** synchronous culture. The data represents an average of three independent biological repeats.

Further, in the synchronous condition, we observed that the basal respiration rate remained constant across all the time points in TOR kinase mutant (**Figure 8C**), similar to the wild-type cells (**Figure 4C**). Upon addition of CCCP, similar to wild-type cells, the OCR levels increased in the light phase but unlike wild-type cells which showed the maximum OCR level at 12 hr of light, TOR kinase mutant showed the maximal OCR level at 6 hr and then it started to decrease reaching a value similar to 0 hr sample (**Figure 8C** and **8E**). These changes indicate that the TOR kinase mutant displayed subdued rhythmicity in mitochondrial activity in the synchronous cultures, which aligns with the reduced rhythmicity observed in case of photosynthetic activity.

Therefore, our study highlights the rhythmic pattern in the chloroplast and mitochondrial activity in the light-dark cycle. This pattern is subdued in case of TOR kinase mutant compared to wild-type cells, suggesting the role of TOR kinase in maintaining the intensity of diurnal changes in the chloroplast and mitochondrial activity.

## Discussion

In photosynthetic organisms, chloroplast and mitochondria play a crucial role in energy metabolism, and their functions are closely interconnected. These organelles dynamically adapt to changing light conditions to maintain cellular homeostasis. Given the critical importance of this organellar cooperation, our study provides the first insight into the coordinated rhythmicity in photosynthetic and mitochondrial activities in *C. reinhardtii* in the 12:12 hr light-dark cycle.

We examined the phenotypic changes including the morphology and positioning of mitochondria vis-à-vis chloroplast cup throughout the diurnal cycle in *C. reinhardtii*. The shift in mitochondrial morphology from tubular (in light phase) to fragmented form (in the dark phase) (**Figure 1A** and **1B**) is consistent with the known light-dark dependent mitochondrial transition from elongated to the spherical form in *C. reinhardtii* (Hiramatsu et al., 2006) and *A. thaliana* (Oikawa et al., 2015). Alongside mitochondrial morphology changes, we documented novel chloroplast morphology shifts in *C. reinhardtii*, from intact cup in light to distortions or puncturing observed in dark (**Figure 1A** and **1B**). Similar 12:12 hr light-dark mediated changes were reported previously in *Acetabularia mediterranea*, where the chloroplast morphology transitions between elongated and spherical forms (Driessche, 1966). Moreover, the increase in the peripheral phenotype of mitochondria vis-à-vis chloroplast cup in the light phase observed in our study (**Figure 1D** and **1E**) aligns with the reported mitochondrial position shifts in *C. reinhardtii* from central to peripheral locations upon transition from high CO_2_ to low/very low CO_2_ conditions, facilitating efficient photosynthesis (Findinier et al., 2024). These coordinated morphological changes resemble physical interactions between chloroplast, mitochondria, and peroxisome in plants, suggesting efficient metabolic cooperation between these organelles (Islam and Takagi, 2010; Oikawa et al., 2015). **Therefore, f**urther we speculated whether this morphological rhythmicity may lead to the rhythmicity in organelle functions.

Expectedly, we observed rhythmicity in both photosynthetic and mitochondrial activity in the diurnal cycle, where both the organelles exhibited peak activity in light phase and a marked decline in the dark (**Figure 9**). Using 77K spectroscopy, we observed that the shift in light-dark phases resulted in altered PSI and PSII activities, that triggered a regulatory mechanism known as state transition, which affected the efficiency and relative abundance of PSI and PSII (**Figure 2C** and **2F**) (Allen, 1992). The relative predominance of PSII over PSI in the light phase suggests a shift to State 1, promoting linear electron flow and thereby enhancing the photosynthetic output (Finazzi et al., 2002). Conversely, in the dark phase, higher PSI over PSII reflects transition to State 2, characterized by active cyclic electron flow and a state of reduced photosynthetic function (Fleischmann et al., 1999; Finazzi et al., 2002). These rhythmic changes in the F (PSI/PSII) ratio were further supported by the corresponding alterations in the protein levels, where we observed elevation of both PsaA and D1 protein levels during the light phase, followed by their decline in the dark phase (**Figure 3C**). The light-dependent regulation of both D1 and PsaA protein has been shown previously, where the synthesis and degradation of the D1 protein is tightly regulated by light intensity to maintain photosynthetic efficiency and prevent photodamage (Aro et al., 1993; Nishiyama et al., 2006) and light-dependent expression of PsaA, reflects its role in optimizing electron transport and energy conversion (Pfannschmidt et al., 1999). These protein level changes and 77K spectral changes (**Figure 2** and **3**) underscore the rhythmic nature of photosynthetic machinery adaptation in the light-dark dependent manner.

**Figure 9:**
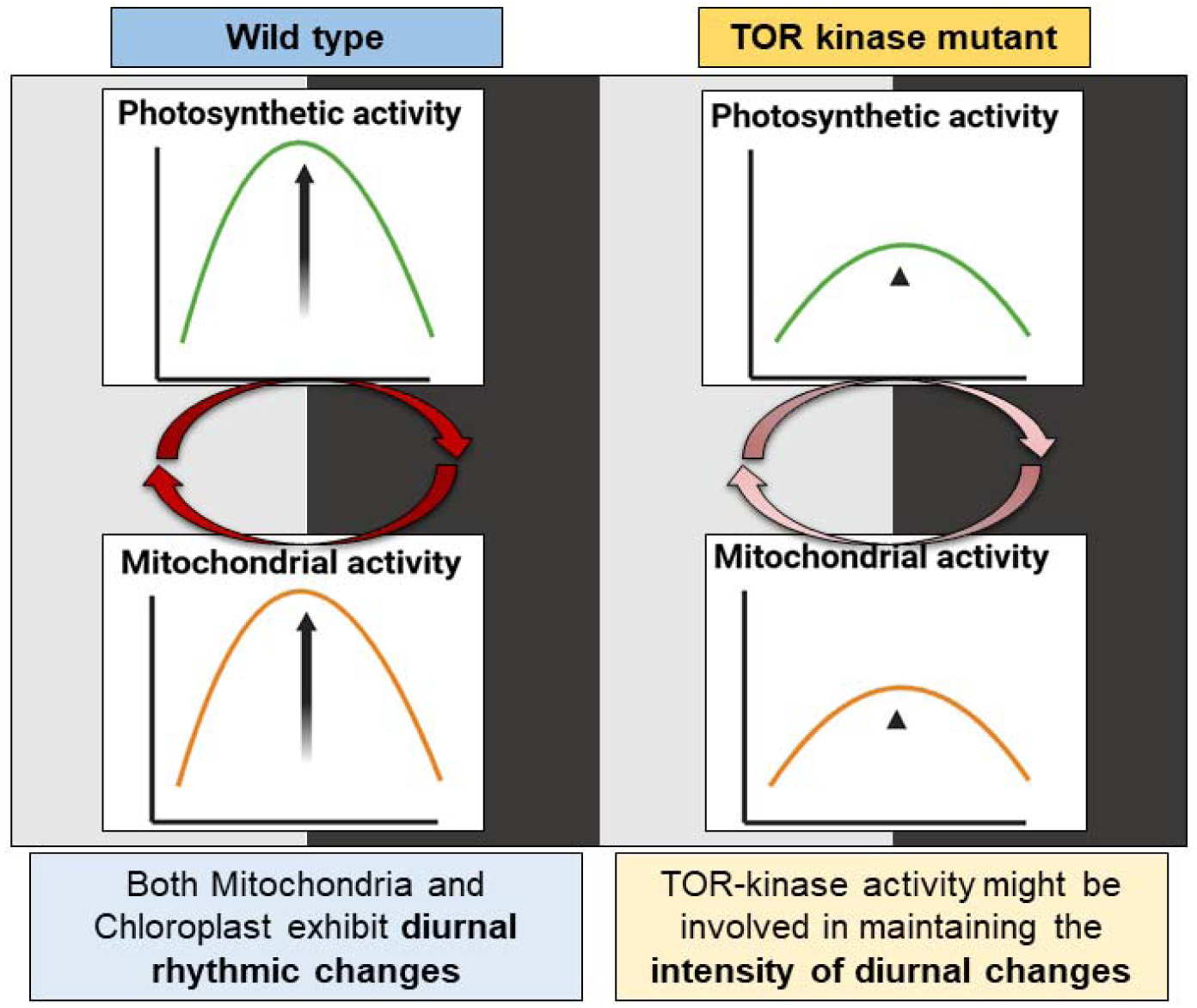
Comparison of photosynthetic and mitochondrial activity changes in *C. reinhardtii* wild-type and TOR kinase mutant in the light-dark cycle. In the wild-type cells, the activity of both chloroplast and mitochondria increases during the day and drops in the dark. There is a rhythmic oscillation in the photosynthetic and mitochondrial activity in the 12:12 hr light/dark dependent manner. TOR kinase mutant also shows similar trend where the activity of both chloroplast and mitochondria increases in the light phase and decreases in the dark phase, but unlike wild-type cells, TOR kinase mutant showed subdued rhythmicity. *Image created using BioRender.com*.

Additionally, we observed an increase in mitochondrial OCR during the light phase compared to the dark (**Figure 4**). Strenkert et al. (2019) has shown similar trend in the mitochondrial respiration capacity in *C. reinhardtii*, suggesting high respiratory activity in the light phase to support ATP demands for metabolism and growth, while reduced activity in the dark phase, which suffices basal cellular requirements. Moreover, in case of higher plants, it is known that the mitochondrial metabolism, particularly the oxidative electron transport and phosphorylation processes, are active in the light condition to maintain the photosynthetic carbon assimilation (Raghavendra and Padmasree, 2003). Altogether, our findings reveal unprecedented coordinated rhythmicity in the chloroplast and mitochondrial functions along with the morphology changes in the diurnal cycle. These rhythmic changes may lead to tight coupling between the two organelles in light phase as opposed to the dark phase, ensuring efficient energy utilization and metabolic flexibility in response to the light-dark cycle (Yoshida and Noguchi, 2011), thus maintaining cellular homeostasis.

A crucial aspect of this homeostasis is the ability to sense and respond to the environmental cues and internal metabolite or nutrient concentration. One of the central and most conserved regulators of these pathways is target of rapamycin (TOR) kinase (Caldana et al., 2019). TOR kinase is essential for detecting a cell’s nutritional and energy state, as a result, it controls the main activities in cells, such as metabolism and cellular respiration (Morita et al., 2013; Dobrenel et al., 2016a). Therefore, we speculated the role of TOR kinase in maintaining the organelle rhythms in the diurnal cycle.

Similar to the control cells, we observed the rhythmicity in mitochondrial and chloroplast morphology upon TOR kinase inhibition (**Figure 5**). However, the extent of rhythmicity was less pronounced in the TOR kinase-inhibited cells. The prevalence of intermediate and fragmented form of mitochondria and distortion in the chloroplast cup upon TOR kinase inhibition relative to the control cells, overlaps with the TOR kinase inhibition mediated fragmentation of mitochondria and the distortions in the chloroplast cup as reported previously (Upadhyaya and Rao, 2019). Additionally, we observed the striking reduction in the peripheral phenotype of mitochondria vis-à-vis chloroplast cup during the light phase of synchrony upon TOR kinase inhibition (**Figure 5C** and **5D**), in contrast to the tight spatial coordination observed in the control cells, which is novel (**Figure 1D** and **1E**). These results suggest subdued rhythmicity in chloroplast and mitochondrial morphology under TOR kinase inhibition compared to the control cells, prompting us to investigate whether the photosynthetic and respiratory activities were also attenuated in the diurnal cycle.

The subdued rhythmicity in the 77K spectra observed in case of TOR kinase mutant (**Figure 6C** and **6F**), relative to the wild-type cells (**Figure 2C** and **2F**), is consistent with the previously documented impairment of PSI and PSII efficiency and inhibition of state transition in response to light availability upon TOR kinase inhibition in *C. reinhardtii* (Upadhyaya and Rao, 2019). Corresponding protein level alterations observed in our study (**Figure 7**) further confirmed the role of TOR kinase in maintaining the photosynthetic rhythmicity across the diurnal cycle. Previously, TOR inactivation in *Arabidopsis* has shown to reduce the photosynthetic protein expression by transcriptional and posttranscriptional downregulation of genes encoding photosynthetic proteins (Dong et al., 2015; Dobrenel et al., 2016b; D’Alessandro et al., 2024) and biosynthetic pathways such as Calvin Benson Cycle proteins (Mallén-Ponce et al., 2022), thereby suggesting the role of TOR kinase in the regulation of photosynthetic proteins. Further, the subdued fluctuations in the OCR values in the TOR kinase mutant (**Figure 8C** and **8E**) in comparison to the wild-type cells (**Figure 4C** and **4E**), paralleled the reduced mitochondrial respiration rate in mammals upon TOR kinase inhibition using rapamycin (Schieke et al., 2006; Ramanathan and Schreiber, 2009). Together, these findings suggest that TOR kinase governs the organellar functions in response to altering light-dark cycle, which aligns with its role in integrating the external light/dark cues with the internal metabolic adjustment to maintain cellular energy homeostasis.

In conclusion, our study highlights the coordination in the rhythmicity of photosynthetic and mitochondrial activities in the light-dark cycle, suggesting the intricate coupling between the two organelles in the light/dark-dependent manner (**Figure 9**). Additionally, we underscore the crucial role of TOR kinase in maintaining the intensity of light-dark dependent rhythmic changes in chloroplast and mitochondrial functions (**Figure 9**). Here, we did not probe for the molecular mechanisms, instead this is the first line of study which led us to speculate the following: (1) Whether the external circadian clock of light-dark cycle is not getting integrated with the internal clock in the TOR kinase mutant and therefore it is leading to the less pronounced or subdued rhythmicity? Intriguingly, TOR kinase depletion or reduction lengthens the circadian period in plants (Urrea-Castellanos et al., 2022), thereby linking TOR kinase signaling to circadian control. (2) TOR kinase may either be a component of central oscillator governing the output pathways, or clock components may be the target of this kinase, as suggested by Mittag et al., 2005. In order to understand the exact role of TOR kinase in circadian rhythms, the system needs to be probed further for molecular interactions, metabolic rhythmicity, and signaling pathways.

## Materials and methods

### Strains, media, and growth condition

*C. reinhardtii* wild-type 5325, TOR kinase mutant 3031 (LMJ.RY0402.203031), and MDH4-GFP (CSI_FC2A03) strains were obtained from the Chlamydomonas Library Project (CLiP) (https://www.chlamylibrary.org), Chlamydomonas resource center (https://www.chlamycollection.org). Cultures were maintained on TAP agar plates (1.5% w/v) as described (Harris, 2009). *C. reinhardtii* strains were grown photoautotrophically at a light intensity of 60 µmol/m²s at 24lC. The light intensity was measured using Model MQ-500 light meter from apogee instruments. We inoculated *C. reinhardtii* colonies from TAP plates into TP liquid medium and setup three different conditions - 1. continuous light condition for 4-5 days (LL); 2. continuous dark condition for 4-5 days (DD); 3. 12:12 hr light: dark cycle (LD) for 4-5 days (by this time, cells reached mid-log-phase and got synchronized) and these cultures were used for further analysis (Bernstein, 1960). All the analyses were performed on samples withdrawn at six hourly intervals for LL, DD, and LD cultures, as specified. In TOR inhibition experiments, AZD8055 (1 µM) (Axon medchem, Axon 1561) was used as TOR kinase inhibitor.

### Confocal imaging for mitochondrial and chloroplast morphology

Live cell confocal imaging was performed using mitochondrial GFP line (MDH4-GFP) for the visualization of mitochondrial morphologies of *C. reinhardtii* control cells and TOR-inhibited cells. For the study of chloroplast morphology, the chloroplast autofluorescence was utilized to observe the changes in the chloroplast. Briefly, the cells (1×10l cells/ml) were immobilized using 0.8% low melting agarose on a coverslip, followed by imaging for mitochondrial GFP (excitation 488 nm; emission 518-574 nm) and autofluorescence of chloroplast (excitation 561 nm; emission 685-750 nm) using appropriate filters in Leica TCS SP8 confocal laser scanning microscope. The images were obtained as Z-stacks and processed with Fiji Software. Mitochondrial morphology was classified as tubular, intermediate, and fragmented, as per the published protocol (Rambold et al., 2011). Each cell was assessed for the mitochondrial phenotype (dominant), and the frequency distribution of cells exhibiting these phenotypes was quantified for untreated and AZD-treated samples. The data from ∼100 cells was statistically analyzed, quantified using ImageJ software, and plotted using GraphPad Prism 9.5.1 software.

### 3D rendering for mitochondria apposition chloroplast

The Z-stack images obtained from the confocal microscopy were 3D reconstructed using Leica TCS SP8 software. By clipping x, y, or z axis, we were able to identify the mitochondria-chloroplast association (Moriyama et al., 2018).

### 77K Spectroscopy

The low temperature (77K) allows trapping of photosynthetic complexes in a specific state, enabling detailed spectroscopic analyses. The fluorescence emission spectrum at 77K was recorded using Jasco FP8500 spectrophotometer. A special dewar flask made of quartz was used to measure the fluorescence at very low temperature (liquid nitrogen temperature). The parameters were set as follows: excitation wavelength 440 nm; emission bandwidth 660-760 nm; excitation and emission slit width 5 nm each. Wild-type and TOR kinase mutant (3×10l cells) at different culture conditions (LL, DD, and LD) were pelleted. The pellets were resuspended in 500 µl TP media, and the cells were immediately taken for fluorescence measurement in the dewar flask (Cardol et al., 2003; Snellenburg et al., 2017). The spectral readings were taken at 0, 6, 12, 18, and 24 hr for each of the conditions. The emission traces were obtained with noise, and the final spectral traces were normalized and plotted using GraphPad prism 9.5.1 Software.

### Western Blotting

Wild-type and TOR kinase mutant cells were harvested at 0, 6, 12, 18, and 24 hr from LL, DD, and LD condition. Total protein was extracted as per the standard protocol and estimated with a BCA assay kit (Thermo Fischer Scientific) using BSA (2 mg/ml) as standard and measured in SPECTROstar Omega spectrophotometer. Proteins (20 µg/lane) were separated by size using SDS-PAGE and transferred onto a nitrocellulose membrane, followed by overnight incubation with primary antibodies, according to the standard protocol. Primary antibodies used: anti-PsaA (Agrisera, AS06 172) 1:5,000; anti-Tubulin alpha chain (Agrisera, AS10 680) 1:5,000; anti-D1 protein of PSII (Agrisera, AS05 084) 1:5,000. Subsequently, the blots were incubated with secondary antibody solution (anti-rabbit IgG Peroxidase conjugated; Sigma, A9169; 1:10,000) for 1 hr at room temperature with agitation. This was followed by developing the blots using Immobilon Forte Western HRP Substrate (Millipore WBLUF0500), as per the manufacturer’s instructions, and imaged using ChemiDocTM MP imaging system (Bio-Rad). The quantification was performed using ImageJ software, and the graphs were plotted using GraphPad Prism 9.5.1 Software.

### Seahorse XFp Flux Analyzer assay

Wild-type and TOR kinase mutant cells at a concentration of 2×10l, were pelleted and resuspended in 500µl of fresh TP media. For the Sea horse mitochondrial flux assay, these cells were pipette into each well in the 8-well flux analyzer plate, followed by measurement of oxygen consumption rate (OCR) as per the Agilent Guidelines (https://www.agilent.com/cs/library/usermanuals/public/XF_Cell_Mito_Stress_Test_Kit_User_Gu <u>ide.pdf</u>). Two of the wells were left blank for blank subtraction. CCCP (1 µM) (Sigma C2759-100 mg) was injected in Port A of each well of the seahorse cartridge. Each measurement comprised of a calibration cycle of 40 minutes followed by an equilibration cycle of 12 minutes, where the cells were mixed and incubated in the dark for 12 minutes, and then the OCR measurements were recorded at 24lC. OCR measurement consisted of 1 minute mix cycle (to oxygenate the microchamber), 1 minute wait period (allow settling of cells), and 3 minute of measurement period (https://www.agilent.com/cs/library/usermanuals/public/S7894-10000_Rev_B_Wave_2_4_User_Guide.pdf). 3 OCR readings were taken for the basal and maximal respiration each. Statistical analyses and the graph were plotted using GraphPad Prism 9.5.1 Software.

## Statistical analysis

All the statistical analyses, including arithmetic mean, standard deviation (SD), and wherever applicable 2-way ANOVA was performed in all cases. 2-way ANOVA ordinary 2 data sets with Bonferroni test was performed using GraphPad Prism version 9.5.1 for windows, GraphPad Software (www.graphpad.com).

## Acknowledgments

We acknowledge the assistance from IISER Tirupati and University of Hyderabad faculty and staff for carrying out the experiments in this study. A special thanks to Prof. N Prakash Prabhu, UoH, for the help in the 77K spectroscopy and Prof. Naresh Babu Sepuri, UoH, for providing the Sea Horse facility in the department. We acknowledge JC Bose fellowship grant (DST) [10X-217 to B.J. Rao], MHRD intramural funding to IISER Tirupati to B.J. Rao; University of Hyderabad funding to B.J. Rao.

## Author contributions

B.J.R. and G.D. conceived and planned the research. G.D. performed the experiments. G.D. and B.J.R. wrote the manuscript. All authors read and approved the final manuscript.

## Supplementary Data

**Supplementary Figure S1: Phosphorylation levels of CrS6K protein. (A)** Western blot analysis of phosphoCrS6K and CrS6K protein was performed for the control cells and TOR kinase-inhibited cells (using AZD8055). The protein levels were normalized with respect to tubulin. **(B)** The ratio of phosphoCrS6K/CrS6K protein was quantified and plotted for both the control cells and TOR kinase inhibited cells. The data represents an average of two independent biological repeats.

**Supplementary Table S1: Quantification of cells in different mitochondrial morphologies in the light-dark cycle. (A)** Percentage of cells in different mitochondrial morphologies – tubular, intermediate, and fragmented. **(B)** Percentage of cells showing peripheral and diffused phenotype of mitochondria apposition with respect to chloroplast. All values represent mean ± SD, n=3.

**Supplementary Table S2: F (PSI/PSII) ratio in the light-dark cycle.** The ratio of fluorescence of PSI/PSII was quantified for all three conditions: **(A)** continuous light (LL), **(B)** continuous dark (DD), and **(C)** 12:12 hour synchrony (LD) across all the time points using wild-type cells. All values represent mean ± SD, n=3.

**Supplementary Table S3: Quantification of cells in different mitochondrial morphologies in the light-dark cycle upon TOR kinase inhibition. (A)** Percentage of cells in different mitochondrial morphologies – tubular, intermediate, and fragmented. **(B)** Percentage of cells showing peripheral and diffused phenotype of mitochondria apposition with respect to chloroplast. All values represent mean ± SD, n=3.

**Supplementary Table S4: F (PSI/PSII) ratio in the light-dark cycle in TOR kinase mutant.** The ratio of fluorescence of PSI/PSII was quantified for all three conditions: **(A)** continuous light (LL), **(B)** continuous dark (DD), and **(C)** 12:12 hour synchrony (LD) across all the time points using TOR kinase mutant. All values represent mean ± SD, n=3.

**Supplementary Video S1: 3D representation of mitochondria and chloroplast morphology in continuous light condition.** Photoautotrophic *C. reinhardtii* mitochondrial GFP (MDH4-GFP) cells in continuous light were imaged using Leica TCS SP8 confocal laser scanning microscope and 3D projected.

**Supplementary Video S2: 3D representation of mitochondria and chloroplast morphology in continuous dark condition.** Photoautotrophic *C. reinhardtii* mitochondrial GFP (MDH4-GFP) cells in continuous dark were imaged using Leica TCS SP8 confocal laser scanning microscope and 3D projected.

**Supplementary Video S3A-S3E: 3D representation of mitochondria and chloroplast morphology in synchronous culture.** Photoautotrophic *C. reinhardtii* mitochondrial GFP (MDH4-GFP) cells in 12:12 hr light-dark cycle were imaged using Leica TCS SP8 confocal laser scanning microscope and 3D projected.

**Supplementary Video S4: 3D representation of peripheral positioning of mitochondria apposition to chloroplast cup in MDH4-GFP cells.**

**Supplementary Video S5: 3D representation of diffused phenotype of mitochondria apposition to chloroplast cup in MDH4-GFP cells.**

**Supplementary Video S6A-S6E: 3D representation of mitochondria and chloroplast morphology in TOR kinase inhibited synchronous culture. Photoautotrophic *C. reinhardtii* mitochondrial GFP (MDH4-GFP) cells in 12:12 hr light-dark cycle were treated with TOR kinase inhibitor and imaged using Leica TCS SP8 confocal laser scanning microscope and 3D projected.**

**Supplementary Video S7: 3D representation of peripheral positioning of mitochondria apposition to chloroplast cup in TOR kinase inhibited cells.**

**Supplementary Video S8: 3D representation of diffused phenotype of mitochondria apposition to chloroplast cup in TOR kinase inhibited cells.**

