## Supplementary figures, tables and videos for "Light-dark dependent rhythmic changes in chloroplast and mitochondrial activity in *Chlamydomonas reinhardtii*"

### Slide 1
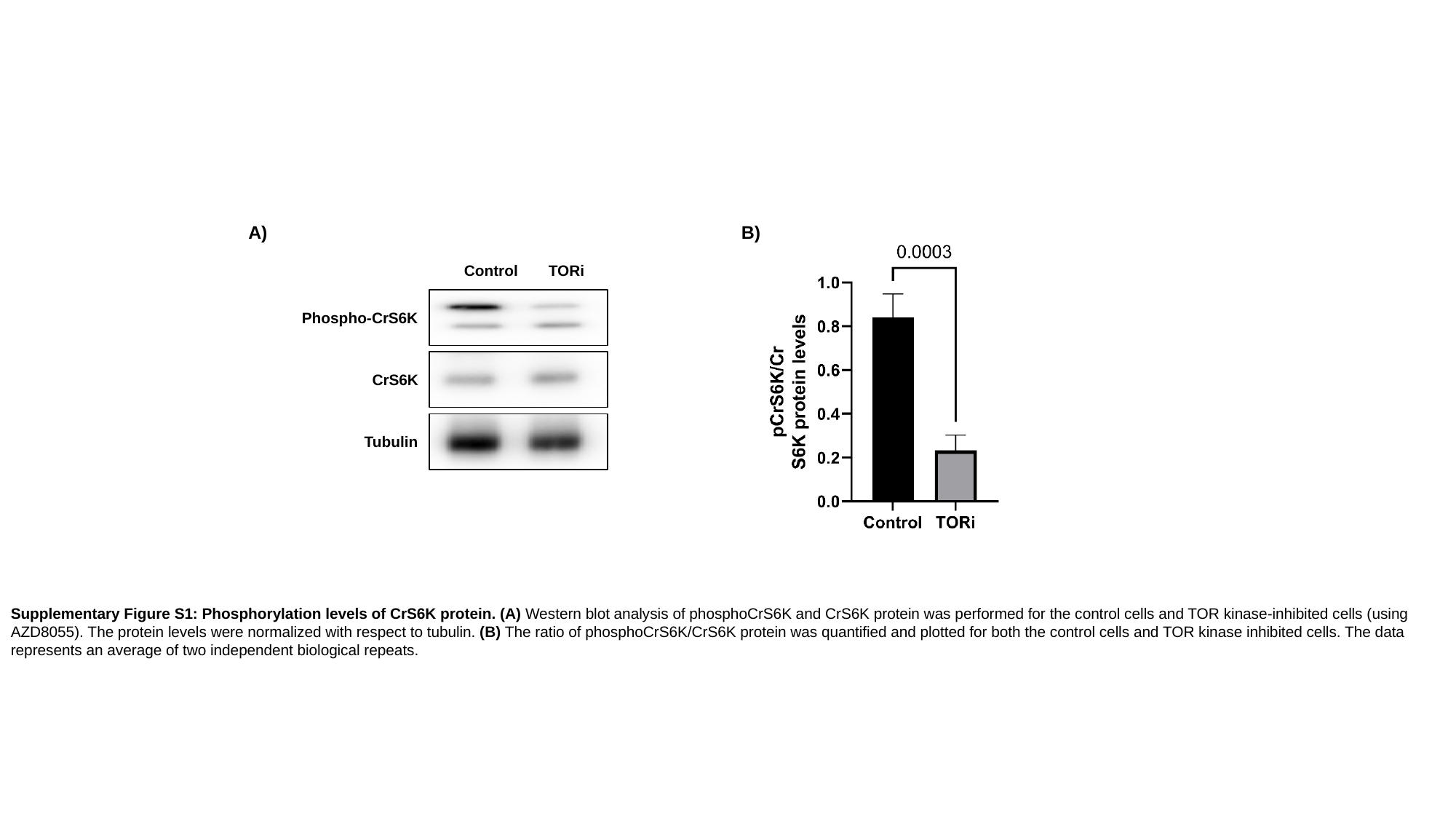

B)
A)
Control
TORi
Phospho-CrS6K
CrS6K
Tubulin
Supplementary Figure S1: Phosphorylation levels of CrS6K protein. (A) Western blot analysis of phosphoCrS6K and CrS6K protein was performed for the control cells and TOR kinase-inhibited cells (using AZD8055). The protein levels were normalized with respect to tubulin. (B) The ratio of phosphoCrS6K/CrS6K protein was quantified and plotted for both the control cells and TOR kinase inhibited cells. The data represents an average of two independent biological repeats.

### Slide 2
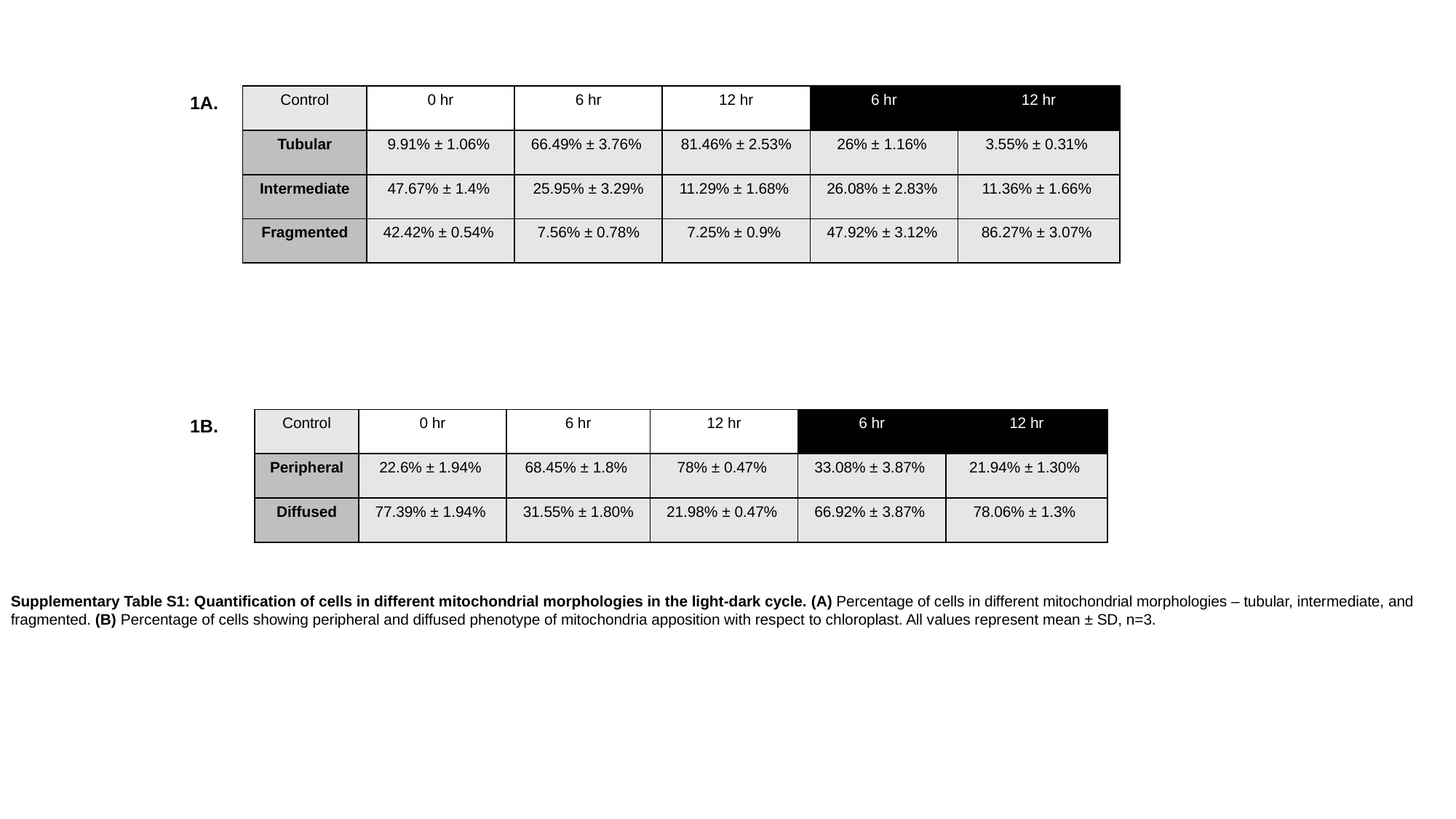

1A.
| Control | 0 hr | 6 hr | 12 hr | 6 hr | 12 hr |
| --- | --- | --- | --- | --- | --- |
| Tubular | 9.91% ± 1.06% | 66.49% ± 3.76% | 81.46% ± 2.53% | 26% ± 1.16% | 3.55% ± 0.31% |
| Intermediate | 47.67% ± 1.4% | 25.95% ± 3.29% | 11.29% ± 1.68% | 26.08% ± 2.83% | 11.36% ± 1.66% |
| Fragmented | 42.42% ± 0.54% | 7.56% ± 0.78% | 7.25% ± 0.9% | 47.92% ± 3.12% | 86.27% ± 3.07% |
1B.
| Control | 0 hr | 6 hr | 12 hr | 6 hr | 12 hr |
| --- | --- | --- | --- | --- | --- |
| Peripheral | 22.6% ± 1.94% | 68.45% ± 1.8% | 78% ± 0.47% | 33.08% ± 3.87% | 21.94% ± 1.30% |
| Diffused | 77.39% ± 1.94% | 31.55% ± 1.80% | 21.98% ± 0.47% | 66.92% ± 3.87% | 78.06% ± 1.3% |
Supplementary Table S1: Quantification of cells in different mitochondrial morphologies in the light-dark cycle. (A) Percentage of cells in different mitochondrial morphologies – tubular, intermediate, and fragmented. (B) Percentage of cells showing peripheral and diffused phenotype of mitochondria apposition with respect to chloroplast. All values represent mean ± SD, n=3.

### Slide 3
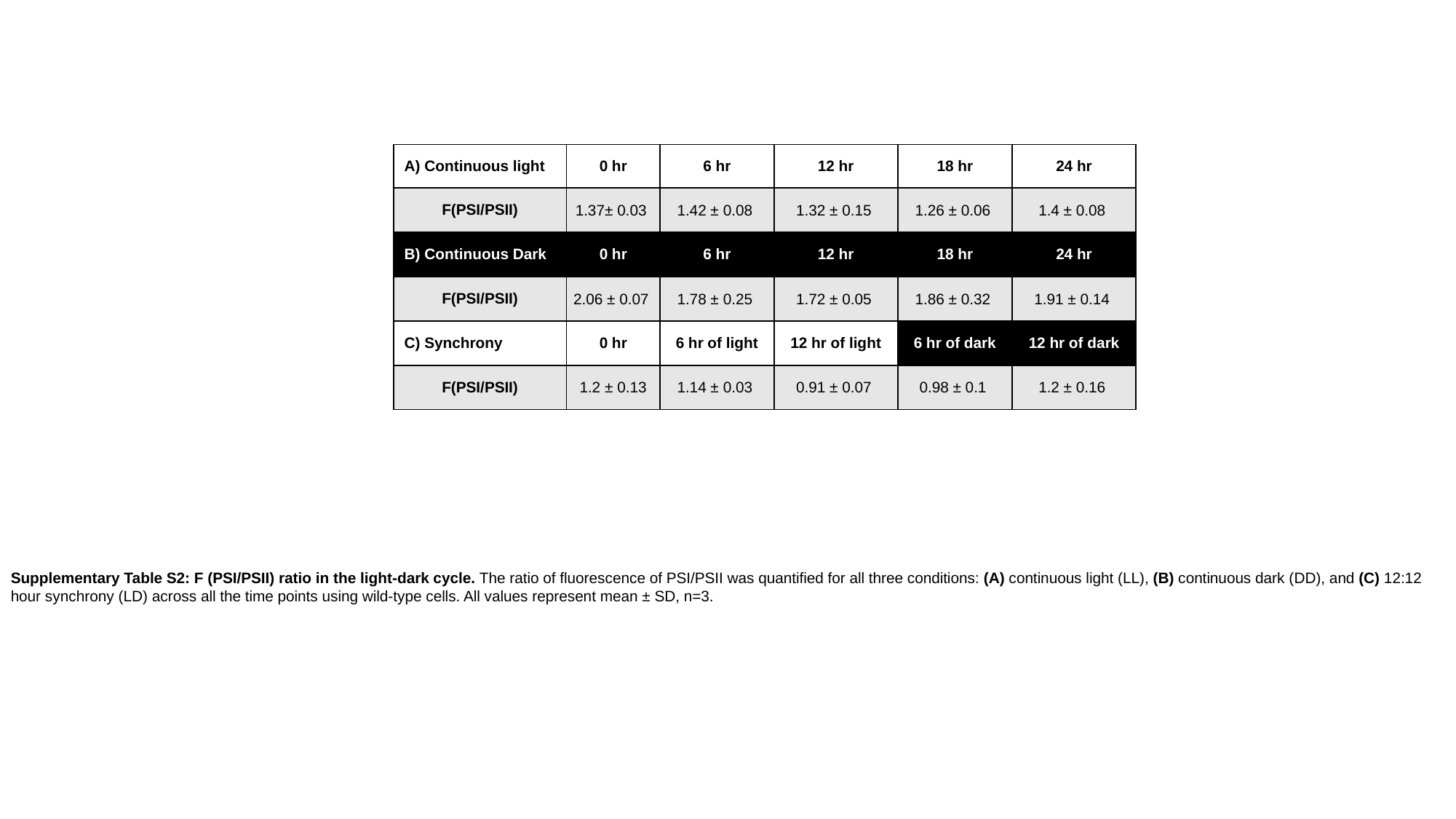

| A) Continuous light | 0 hr | 6 hr | 12 hr | 18 hr | 24 hr |
| --- | --- | --- | --- | --- | --- |
| F(PSI/PSII) | 1.37± 0.03 | 1.42 ± 0.08 | 1.32 ± 0.15 | 1.26 ± 0.06 | 1.4 ± 0.08 |
| B) Continuous Dark | 0 hr | 6 hr | 12 hr | 18 hr | 24 hr |
| F(PSI/PSII) | 2.06 ± 0.07 | 1.78 ± 0.25 | 1.72 ± 0.05 | 1.86 ± 0.32 | 1.91 ± 0.14 |
| C) Synchrony | 0 hr | 6 hr of light | 12 hr of light | 6 hr of dark | 12 hr of dark |
| F(PSI/PSII) | 1.2 ± 0.13 | 1.14 ± 0.03 | 0.91 ± 0.07 | 0.98 ± 0.1 | 1.2 ± 0.16 |
Supplementary Table S2: F (PSI/PSII) ratio in the light-dark cycle. The ratio of fluorescence of PSI/PSII was quantified for all three conditions: (A) continuous light (LL), (B) continuous dark (DD), and (C) 12:12 hour synchrony (LD) across all the time points using wild-type cells. All values represent mean ± SD, n=3.

### Slide 4
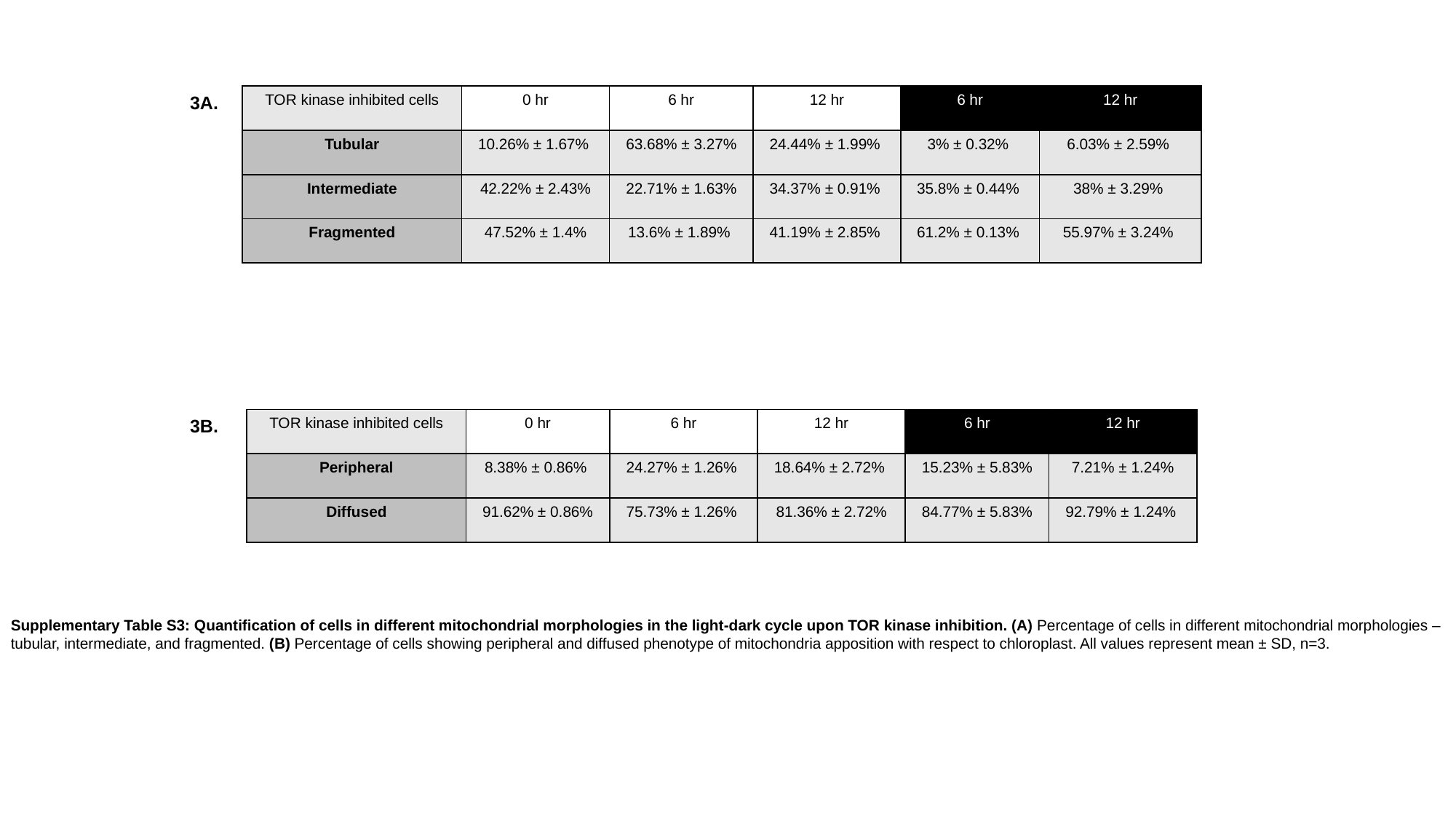

3A.
| TOR kinase inhibited cells | 0 hr | 6 hr | 12 hr | 6 hr | 12 hr |
| --- | --- | --- | --- | --- | --- |
| Tubular | 10.26% ± 1.67% | 63.68% ± 3.27% | 24.44% ± 1.99% | 3% ± 0.32% | 6.03% ± 2.59% |
| Intermediate | 42.22% ± 2.43% | 22.71% ± 1.63% | 34.37% ± 0.91% | 35.8% ± 0.44% | 38% ± 3.29% |
| Fragmented | 47.52% ± 1.4% | 13.6% ± 1.89% | 41.19% ± 2.85% | 61.2% ± 0.13% | 55.97% ± 3.24% |
3B.
| TOR kinase inhibited cells | 0 hr | 6 hr | 12 hr | 6 hr | 12 hr |
| --- | --- | --- | --- | --- | --- |
| Peripheral | 8.38% ± 0.86% | 24.27% ± 1.26% | 18.64% ± 2.72% | 15.23% ± 5.83% | 7.21% ± 1.24% |
| Diffused | 91.62% ± 0.86% | 75.73% ± 1.26% | 81.36% ± 2.72% | 84.77% ± 5.83% | 92.79% ± 1.24% |
Supplementary Table S3: Quantification of cells in different mitochondrial morphologies in the light-dark cycle upon TOR kinase inhibition. (A) Percentage of cells in different mitochondrial morphologies – tubular, intermediate, and fragmented. (B) Percentage of cells showing peripheral and diffused phenotype of mitochondria apposition with respect to chloroplast. All values represent mean ± SD, n=3.

### Slide 5
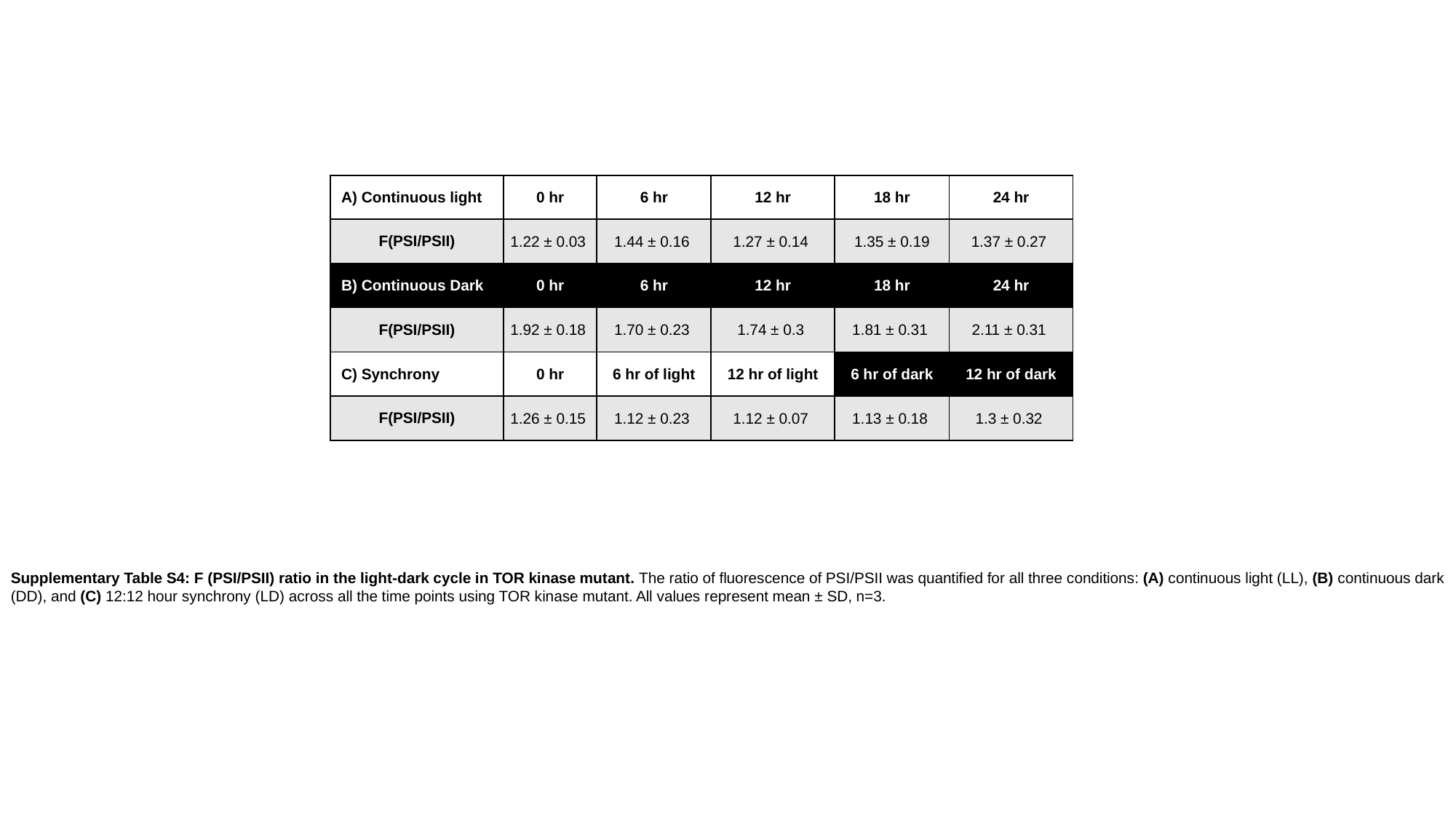

| A) Continuous light | 0 hr | 6 hr | 12 hr | 18 hr | 24 hr |
| --- | --- | --- | --- | --- | --- |
| F(PSI/PSII) | 1.22 ± 0.03 | 1.44 ± 0.16 | 1.27 ± 0.14 | 1.35 ± 0.19 | 1.37 ± 0.27 |
| B) Continuous Dark | 0 hr | 6 hr | 12 hr | 18 hr | 24 hr |
| F(PSI/PSII) | 1.92 ± 0.18 | 1.70 ± 0.23 | 1.74 ± 0.3 | 1.81 ± 0.31 | 2.11 ± 0.31 |
| C) Synchrony | 0 hr | 6 hr of light | 12 hr of light | 6 hr of dark | 12 hr of dark |
| F(PSI/PSII) | 1.26 ± 0.15 | 1.12 ± 0.23 | 1.12 ± 0.07 | 1.13 ± 0.18 | 1.3 ± 0.32 |
Supplementary Table S4: F (PSI/PSII) ratio in the light-dark cycle in TOR kinase mutant. The ratio of fluorescence of PSI/PSII was quantified for all three conditions: (A) continuous light (LL), (B) continuous dark (DD), and (C) 12:12 hour synchrony (LD) across all the time points using TOR kinase mutant. All values represent mean ± SD, n=3.

### Slide 6
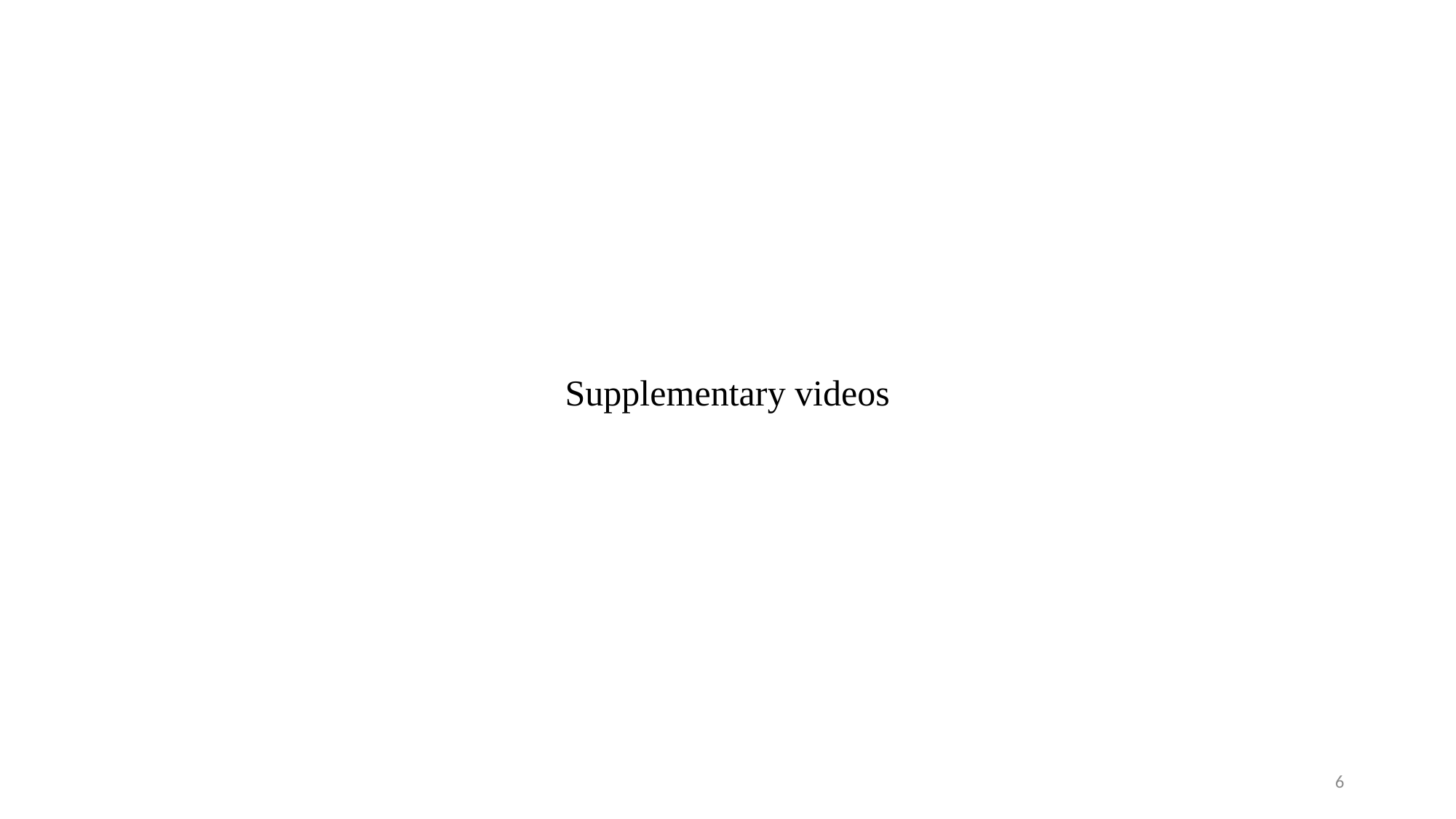

#### Supplementary videos
6

### Slide 7
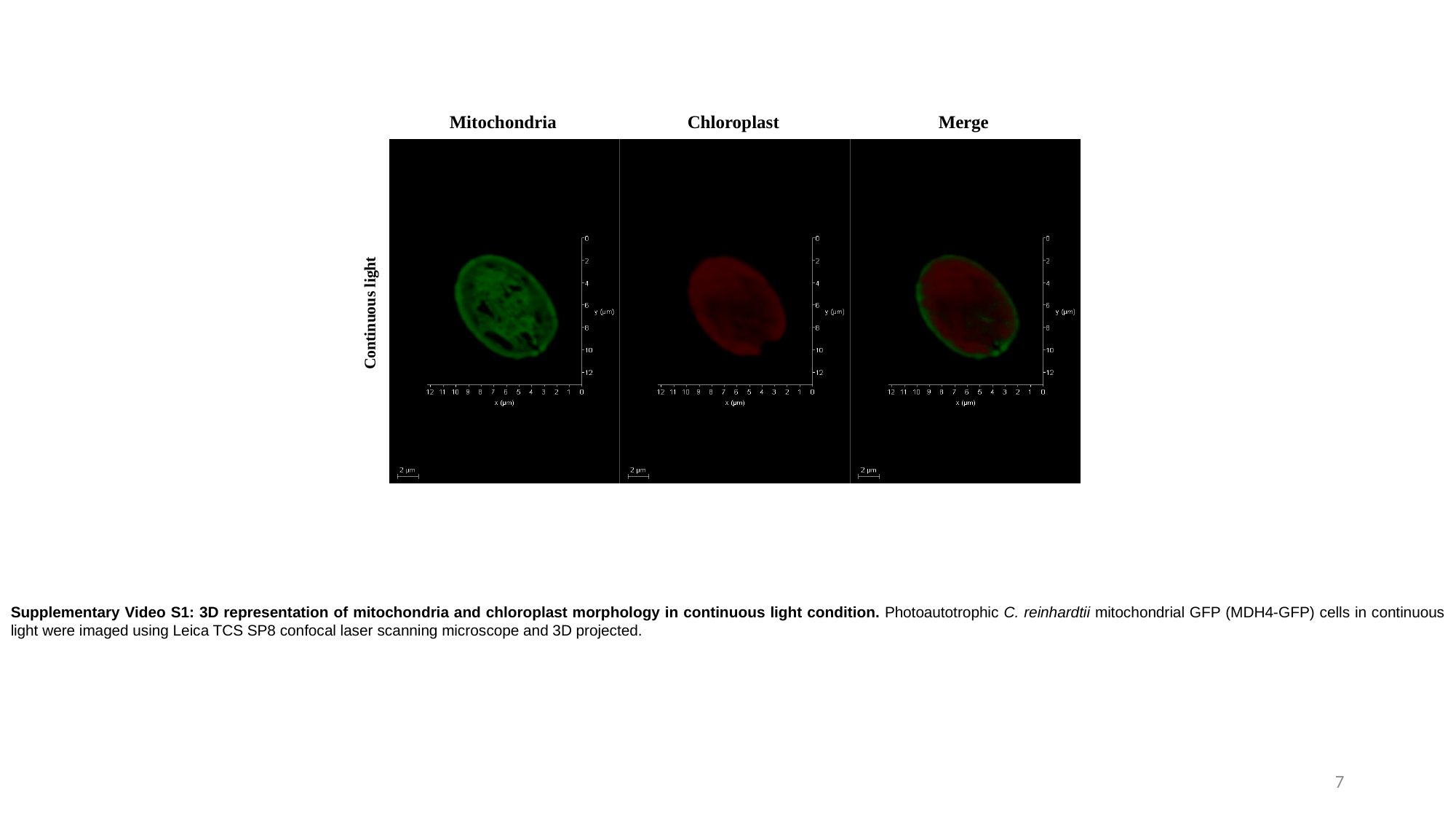

Mitochondria
Chloroplast
Merge
Continuous light
Supplementary Video S1: 3D representation of mitochondria and chloroplast morphology in continuous light condition. Photoautotrophic C. reinhardtii mitochondrial GFP (MDH4-GFP) cells in continuous light were imaged using Leica TCS SP8 confocal laser scanning microscope and 3D projected.
7

### Slide 8
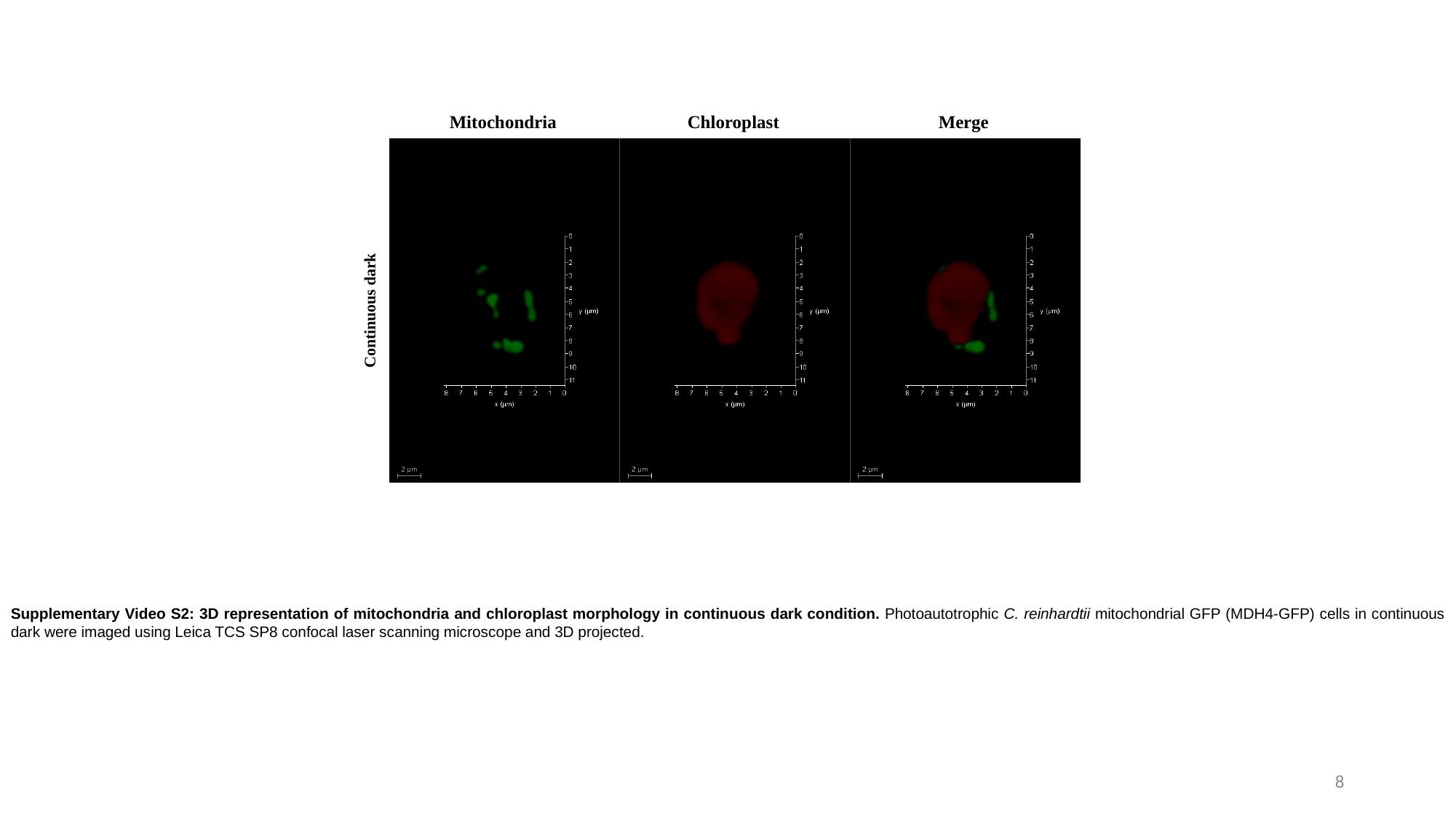

Mitochondria
Chloroplast
Merge
Continuous dark
Supplementary Video S2: 3D representation of mitochondria and chloroplast morphology in continuous dark condition. Photoautotrophic C. reinhardtii mitochondrial GFP (MDH4-GFP) cells in continuous dark were imaged using Leica TCS SP8 confocal laser scanning microscope and 3D projected.
8

### Slide 9
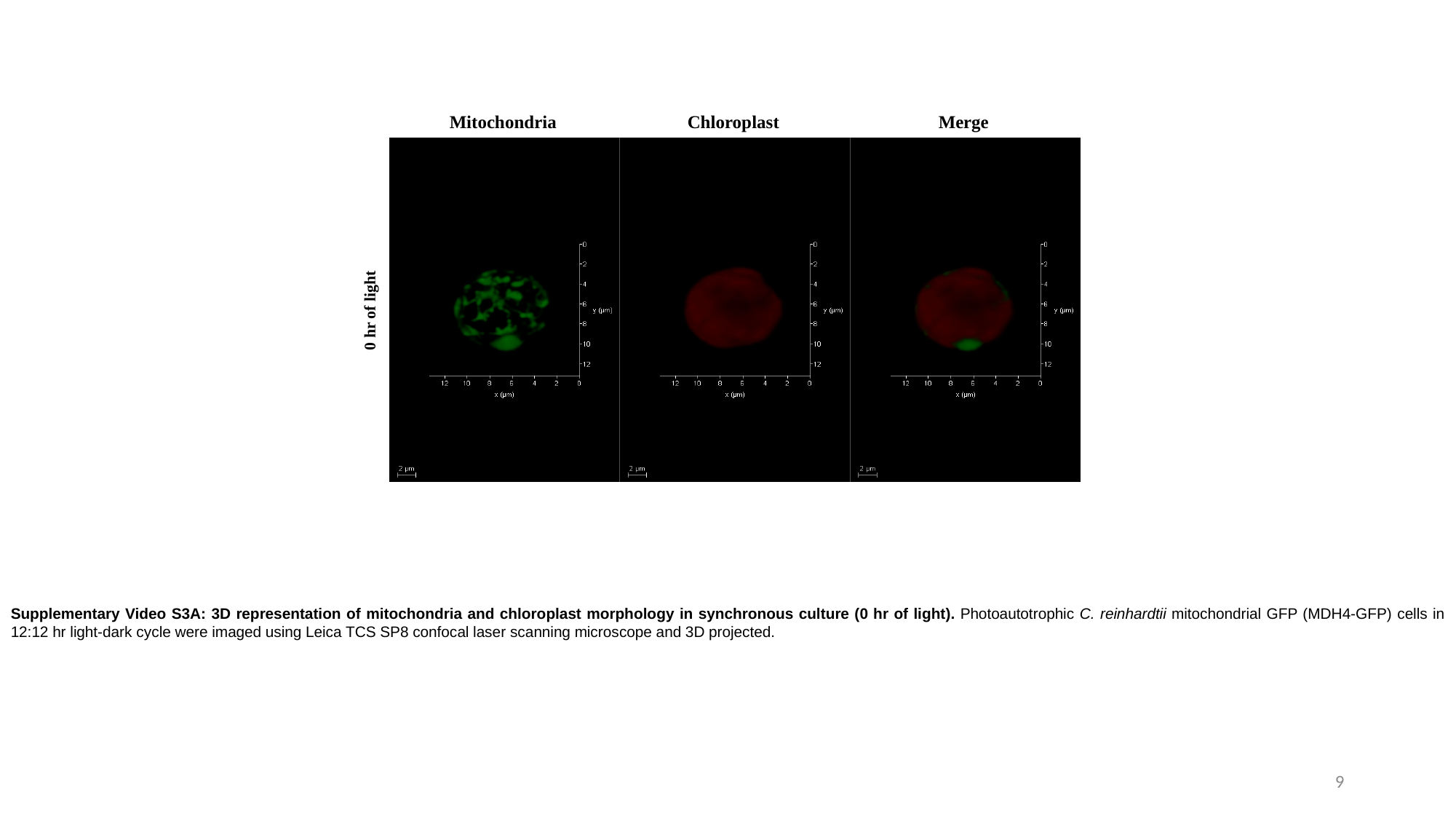

Mitochondria
Chloroplast
Merge
0 hr of light
Supplementary Video S3A: 3D representation of mitochondria and chloroplast morphology in synchronous culture (0 hr of light). Photoautotrophic C. reinhardtii mitochondrial GFP (MDH4-GFP) cells in 12:12 hr light-dark cycle were imaged using Leica TCS SP8 confocal laser scanning microscope and 3D projected.
9

### Slide 10
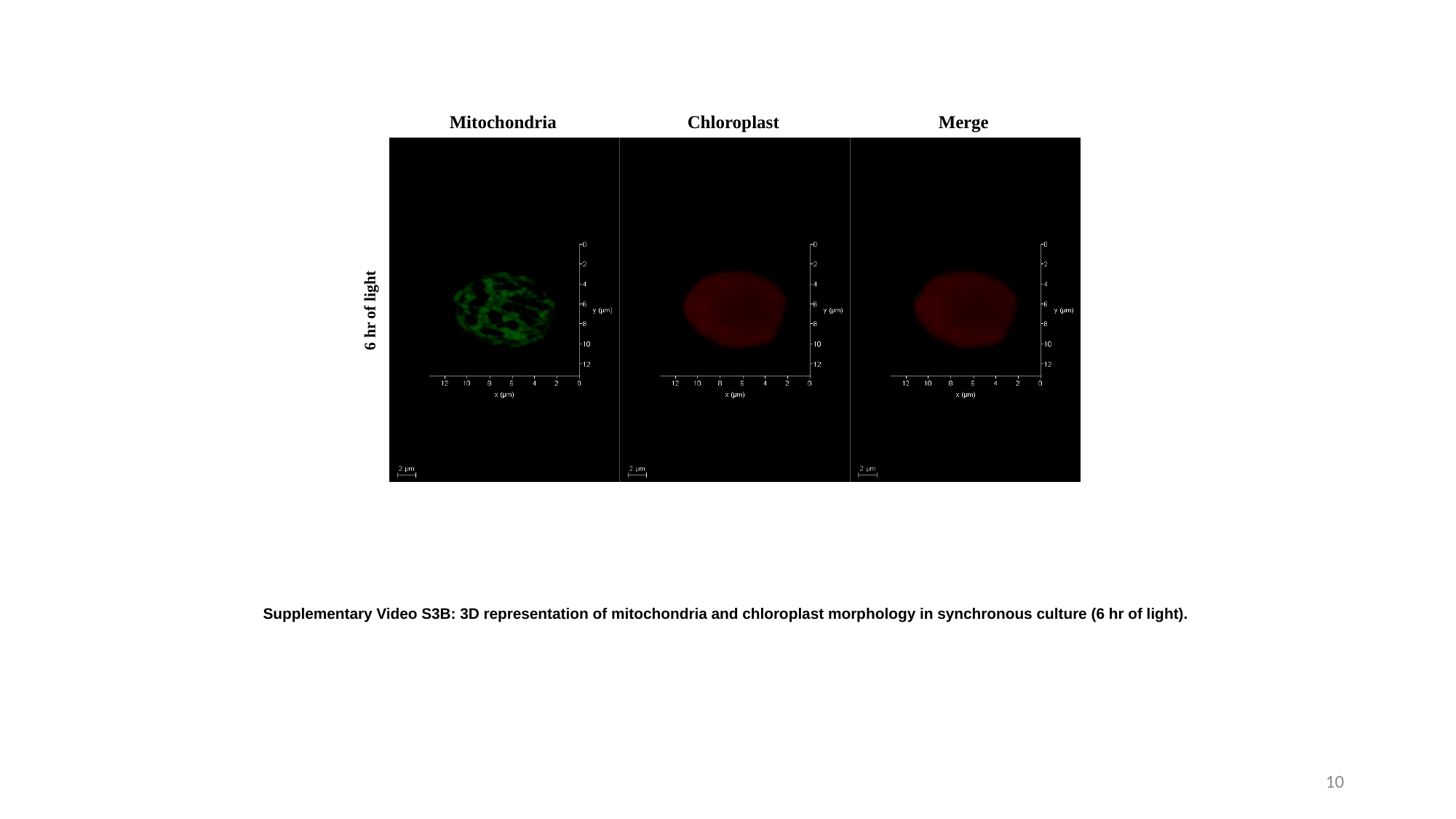

Mitochondria
Chloroplast
Merge
6 hr of light
Supplementary Video S3B: 3D representation of mitochondria and chloroplast morphology in synchronous culture (6 hr of light).
10

### Slide 11
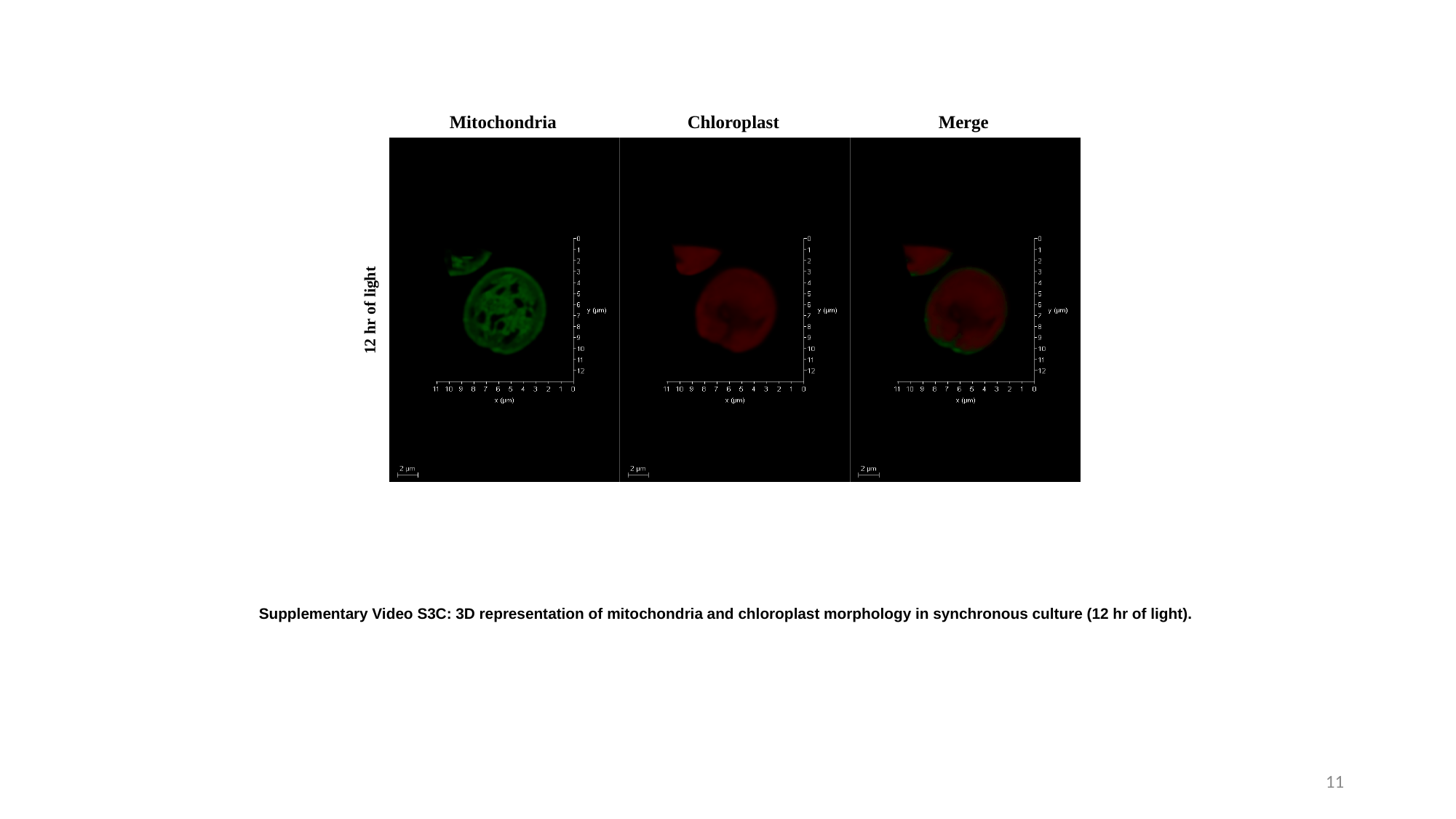

Mitochondria
Chloroplast
Merge
12 hr of light
Supplementary Video S3C: 3D representation of mitochondria and chloroplast morphology in synchronous culture (12 hr of light).
11

### Slide 12
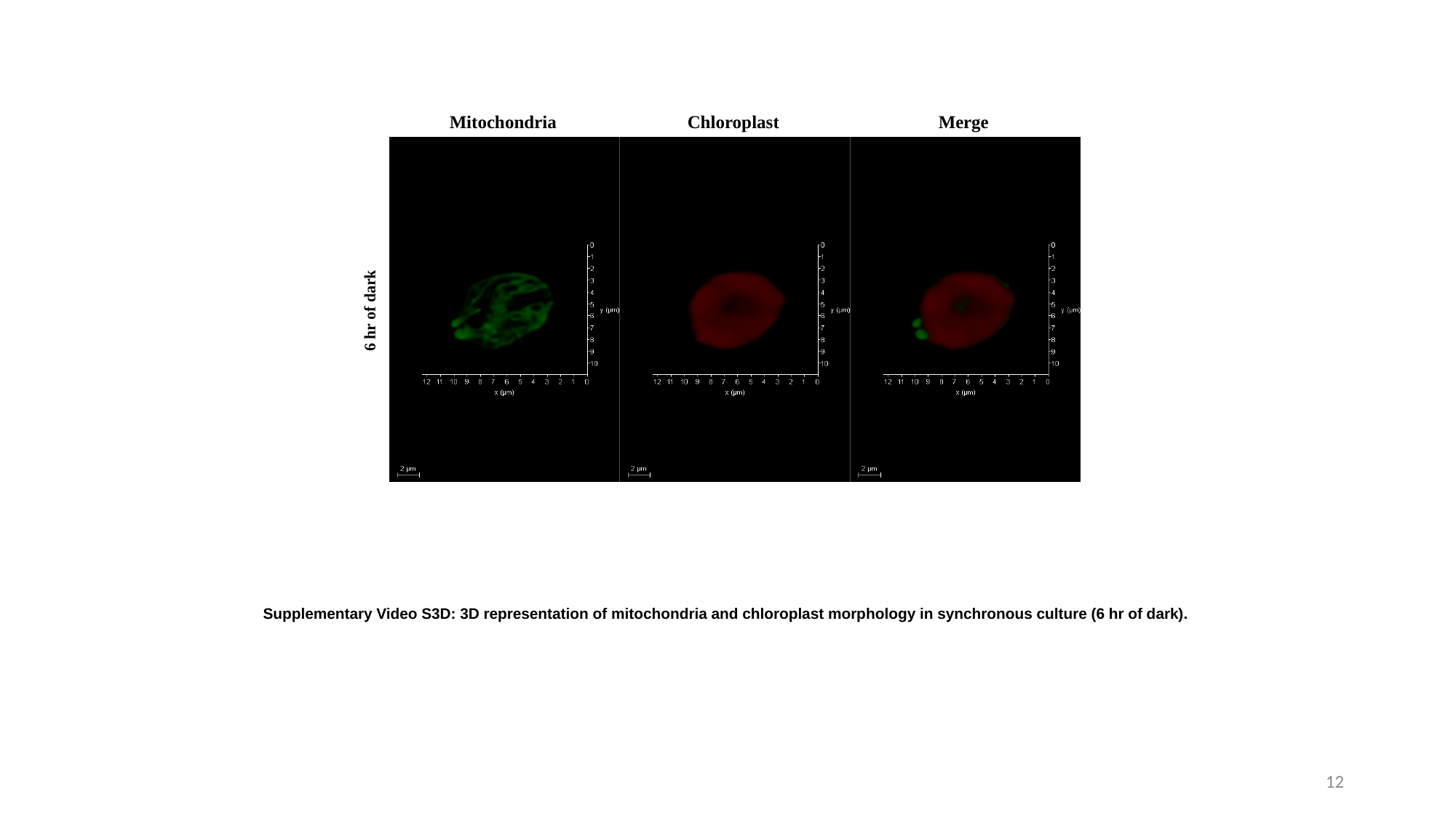

Mitochondria
Chloroplast
Merge
6 hr of dark
Supplementary Video S3D: 3D representation of mitochondria and chloroplast morphology in synchronous culture (6 hr of dark).
12

### Slide 13
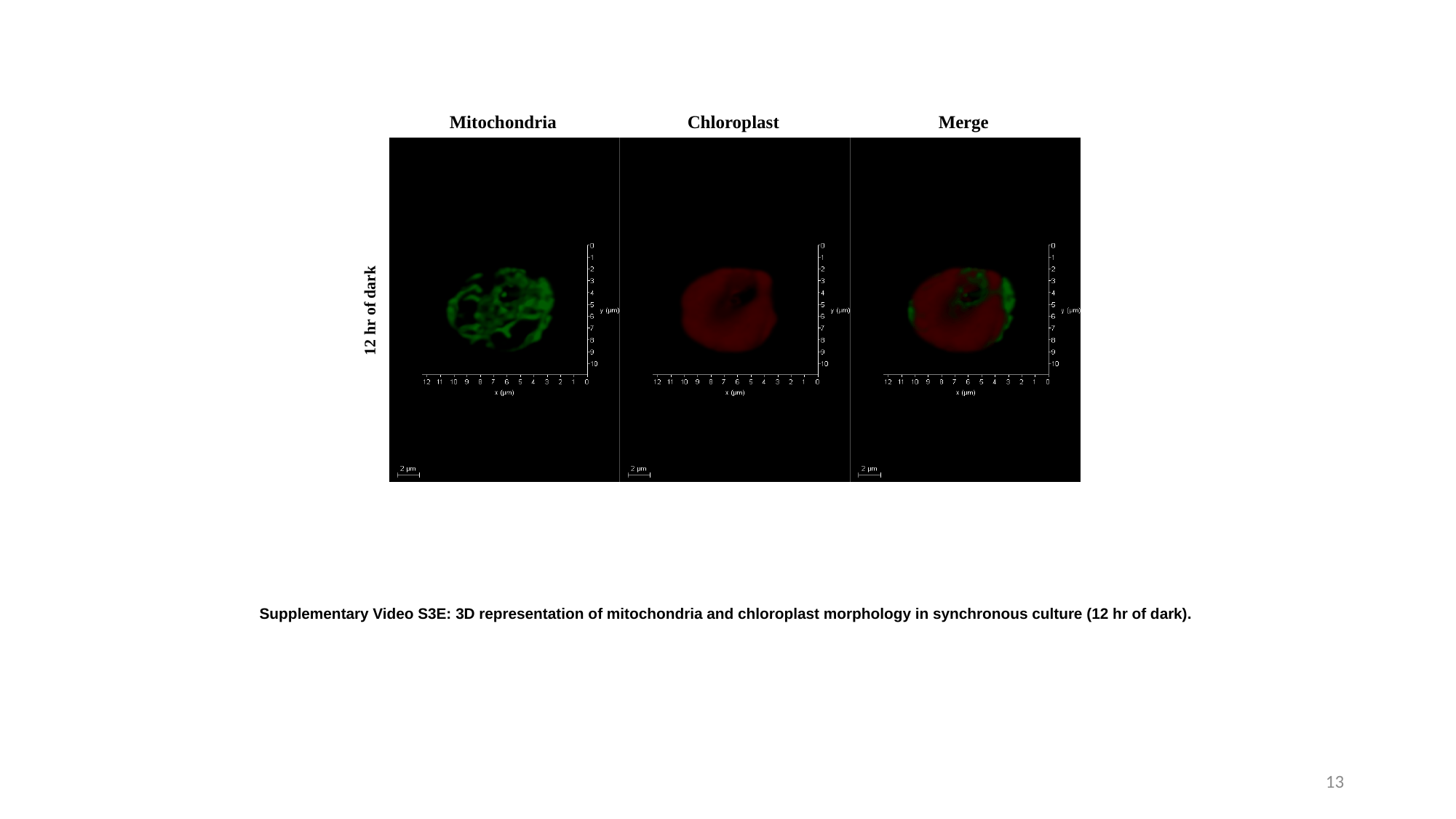

Mitochondria
Chloroplast
Merge
12 hr of dark
Supplementary Video S3E: 3D representation of mitochondria and chloroplast morphology in synchronous culture (12 hr of dark).
13

### Slide 14
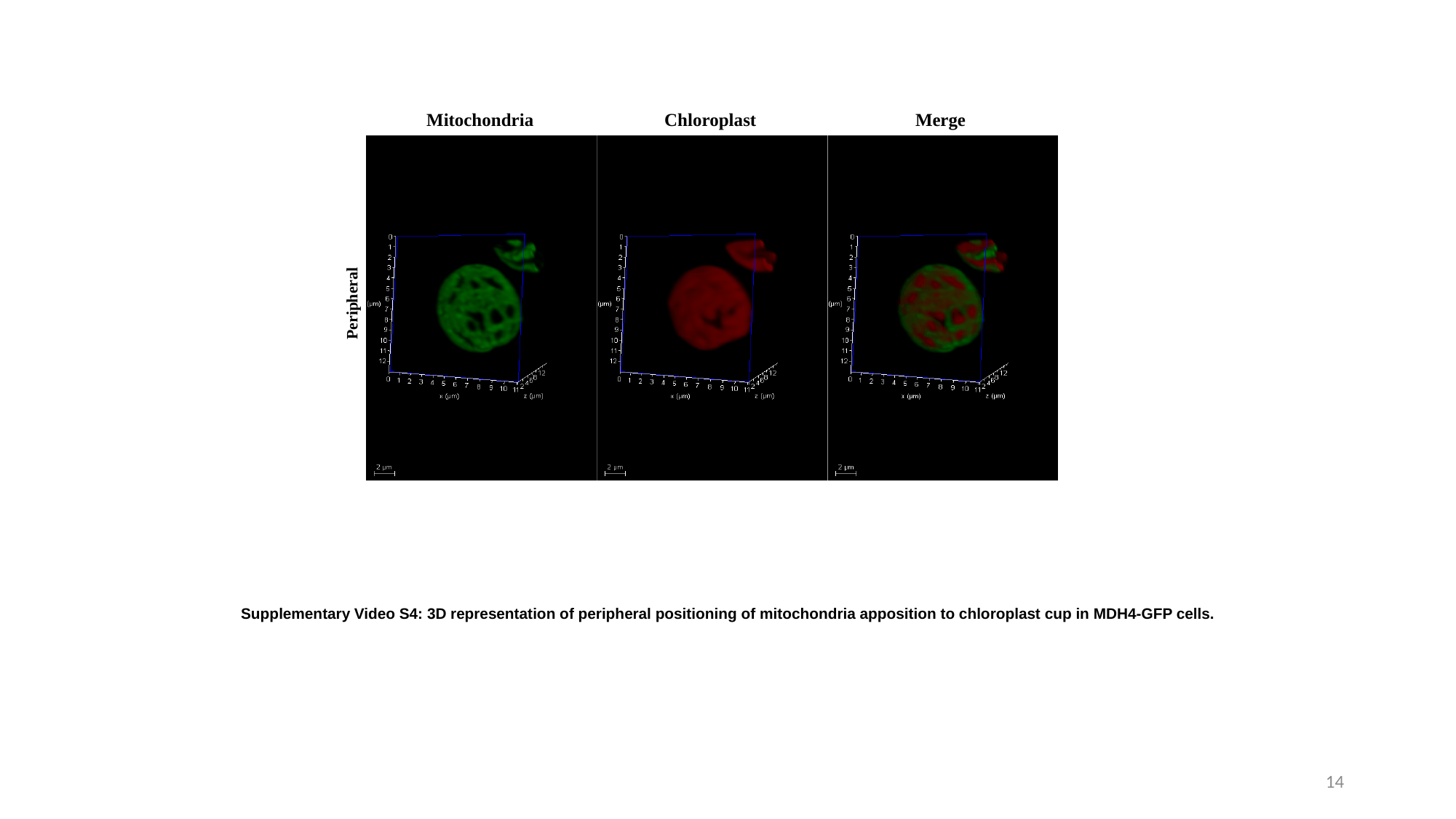

Mitochondria
Chloroplast
Merge
Peripheral
Supplementary Video S4: 3D representation of peripheral positioning of mitochondria apposition to chloroplast cup in MDH4-GFP cells.
14

### Slide 15
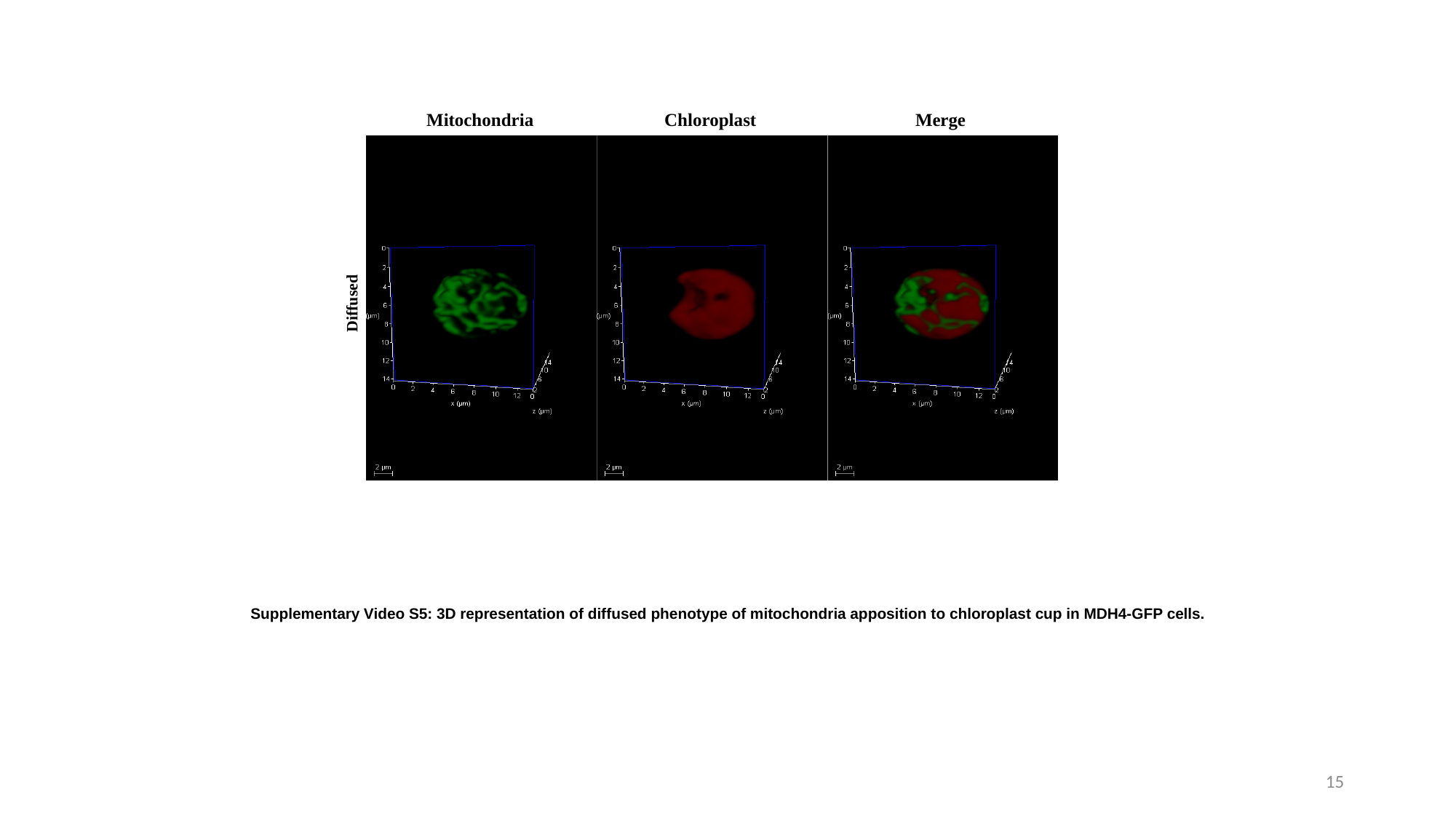

Mitochondria
Chloroplast
Merge
Diffused
Supplementary Video S5: 3D representation of diffused phenotype of mitochondria apposition to chloroplast cup in MDH4-GFP cells.
15

### Slide 16
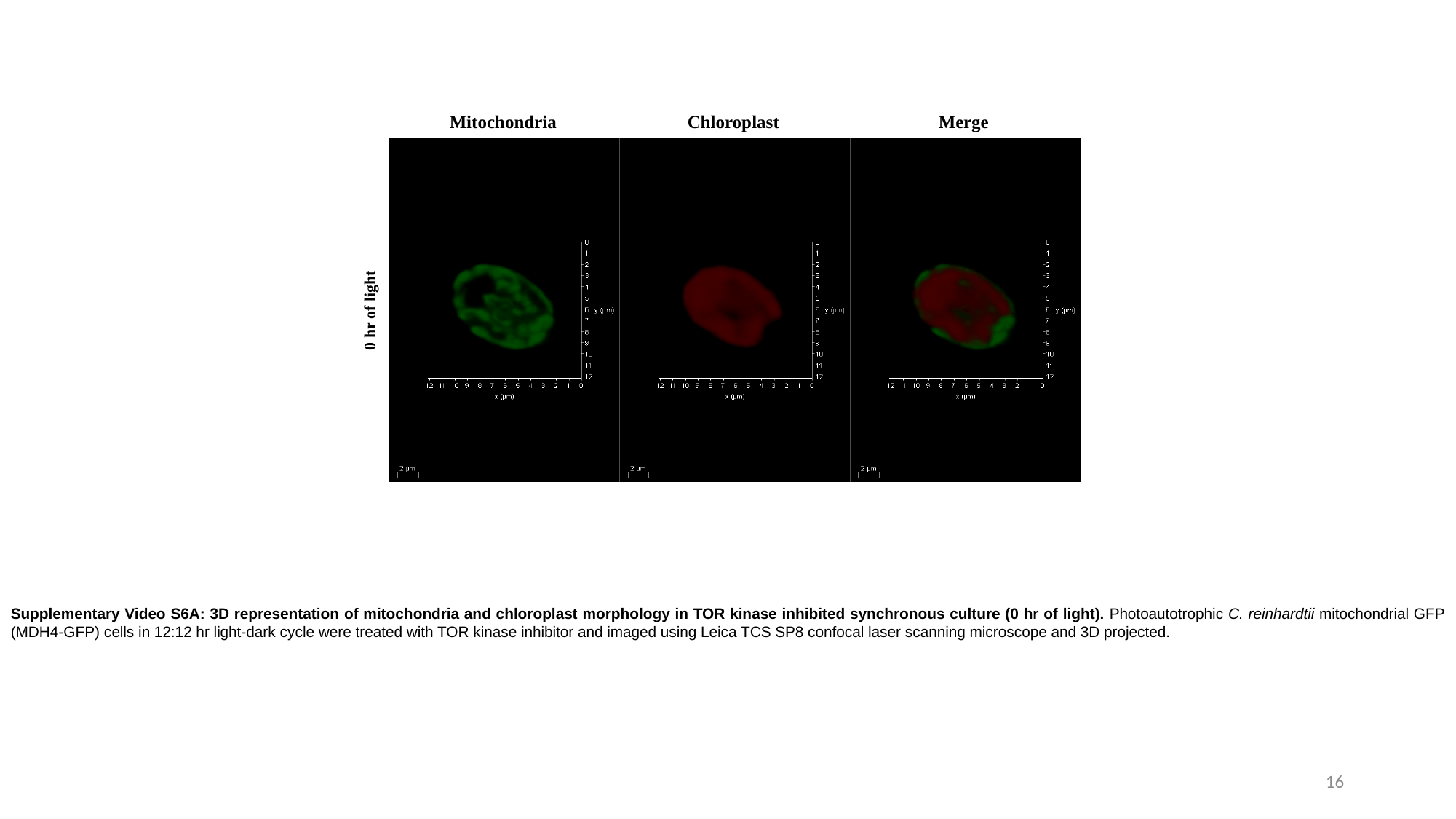

Mitochondria
Chloroplast
Merge
0 hr of light
Supplementary Video S6A: 3D representation of mitochondria and chloroplast morphology in TOR kinase inhibited synchronous culture (0 hr of light). Photoautotrophic C. reinhardtii mitochondrial GFP (MDH4-GFP) cells in 12:12 hr light-dark cycle were treated with TOR kinase inhibitor and imaged using Leica TCS SP8 confocal laser scanning microscope and 3D projected.
16

### Slide 17
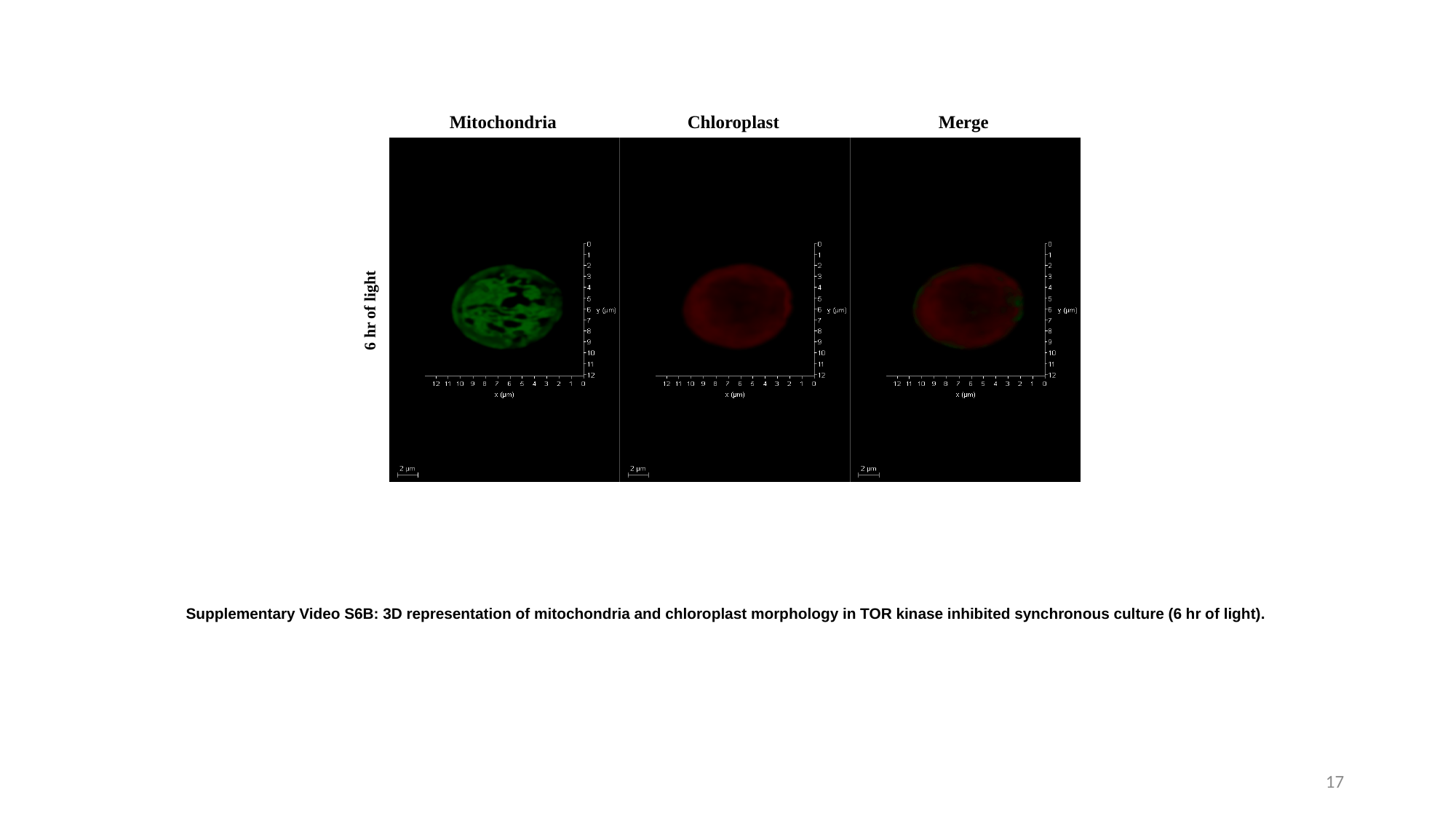

Mitochondria
Chloroplast
Merge
6 hr of light
Supplementary Video S6B: 3D representation of mitochondria and chloroplast morphology in TOR kinase inhibited synchronous culture (6 hr of light).
17

### Slide 18
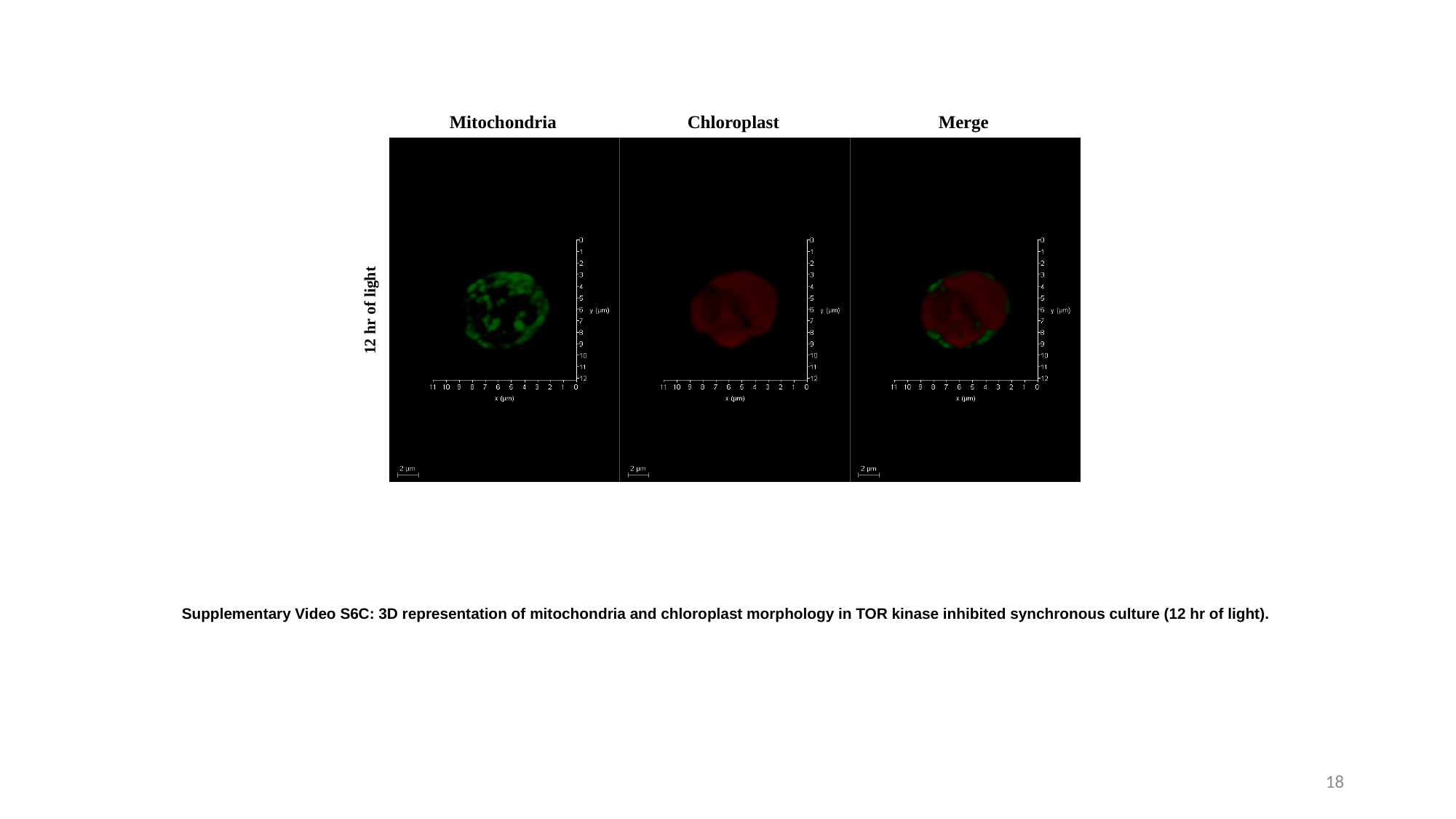

Mitochondria
Chloroplast
Merge
12 hr of light
Supplementary Video S6C: 3D representation of mitochondria and chloroplast morphology in TOR kinase inhibited synchronous culture (12 hr of light).
18

### Slide 19
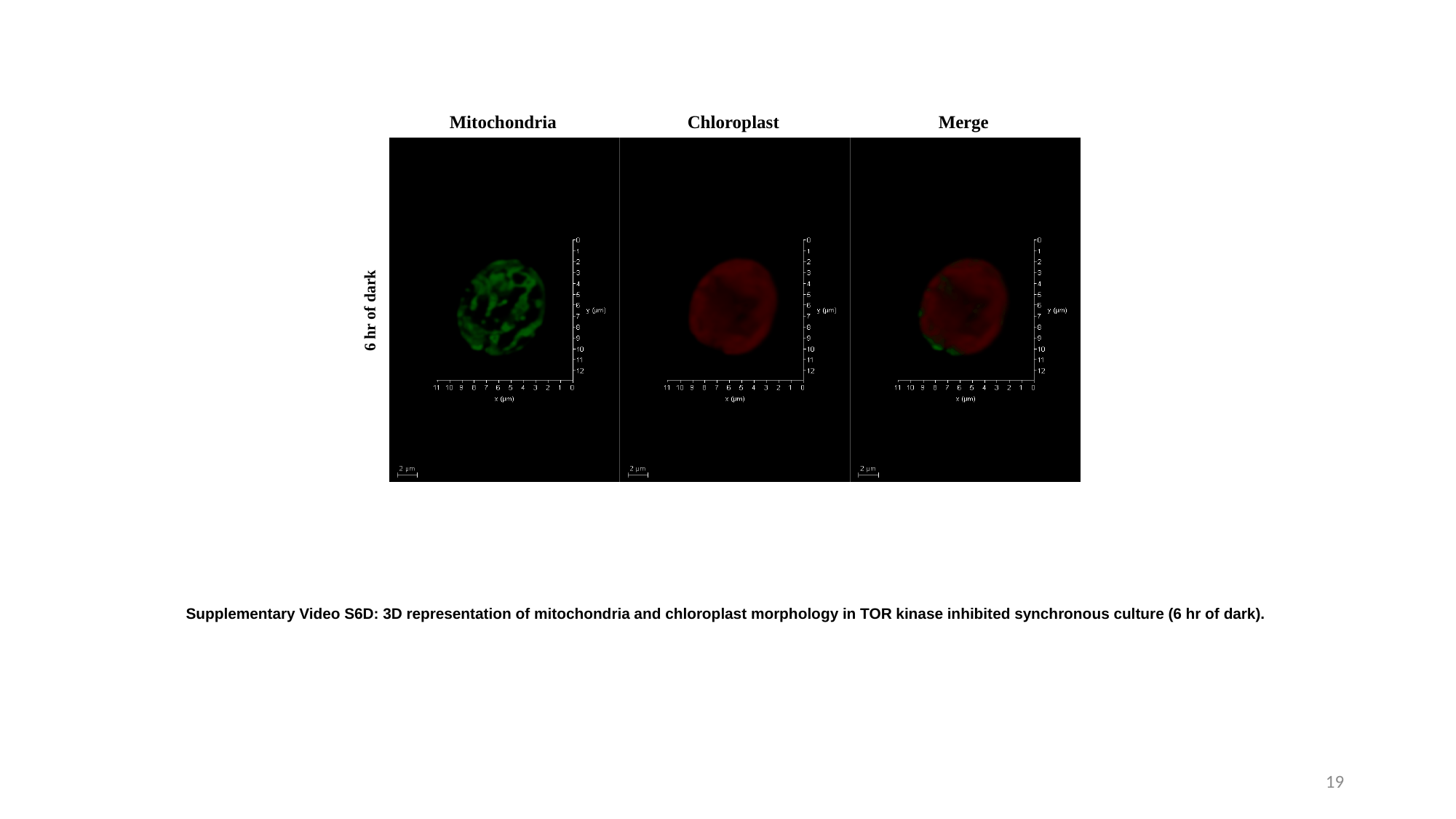

Mitochondria
Chloroplast
Merge
6 hr of dark
Supplementary Video S6D: 3D representation of mitochondria and chloroplast morphology in TOR kinase inhibited synchronous culture (6 hr of dark).
19

### Slide 20
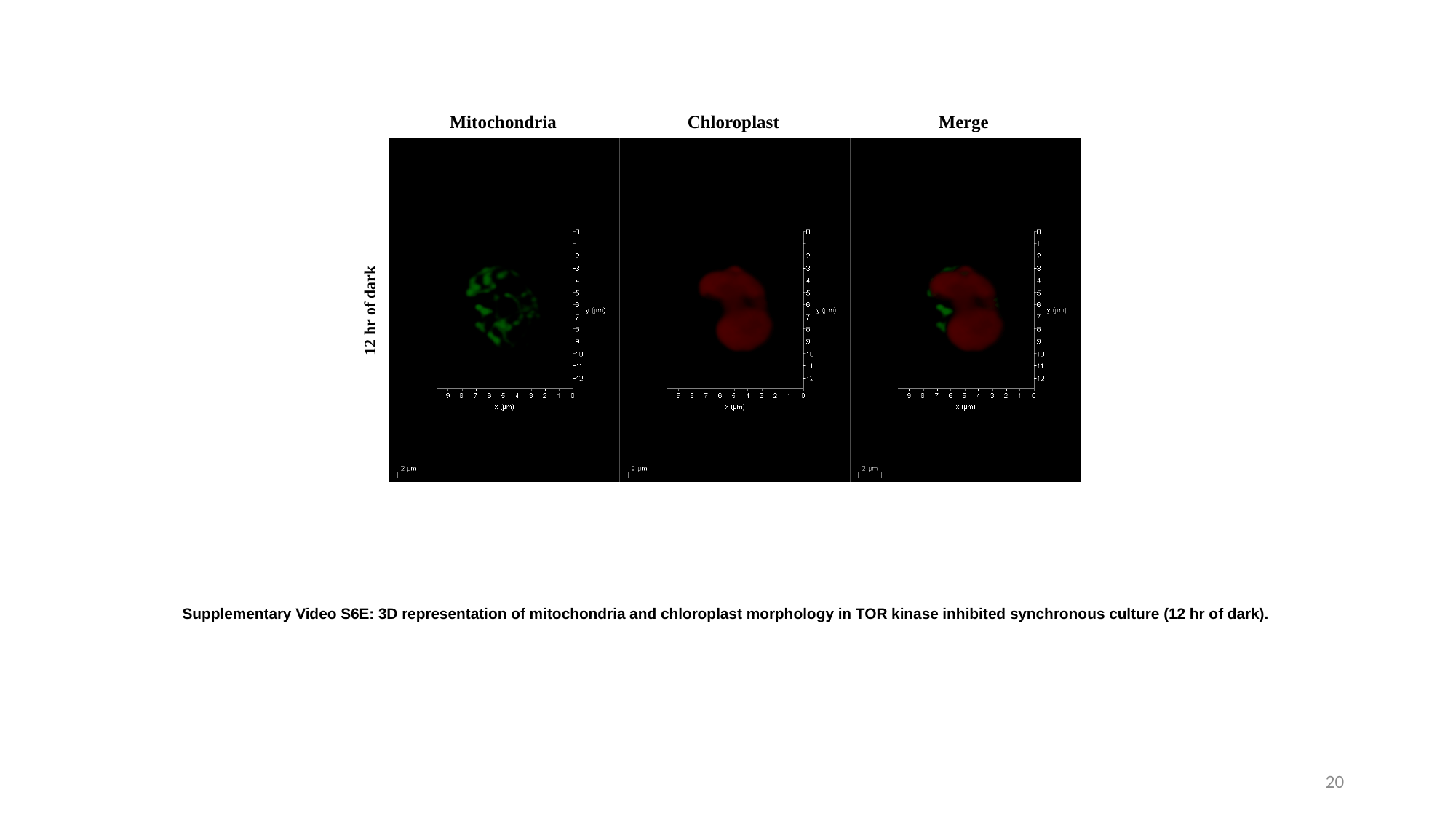

Mitochondria
Chloroplast
Merge
12 hr of dark
Supplementary Video S6E: 3D representation of mitochondria and chloroplast morphology in TOR kinase inhibited synchronous culture (12 hr of dark).
20

### Slide 21
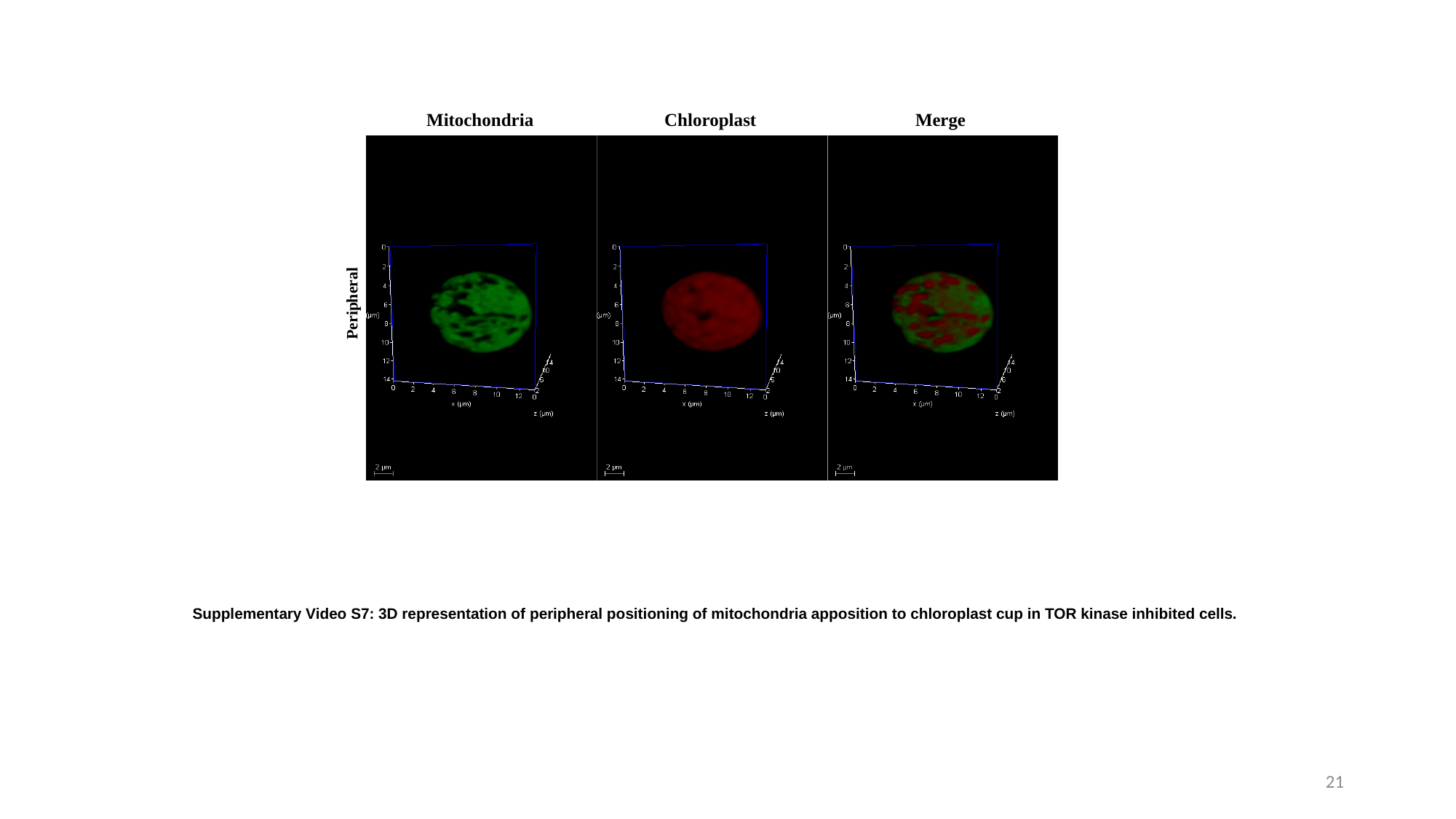

Mitochondria
Chloroplast
Merge
Peripheral
Supplementary Video S7: 3D representation of peripheral positioning of mitochondria apposition to chloroplast cup in TOR kinase inhibited cells.
21

### Slide 22
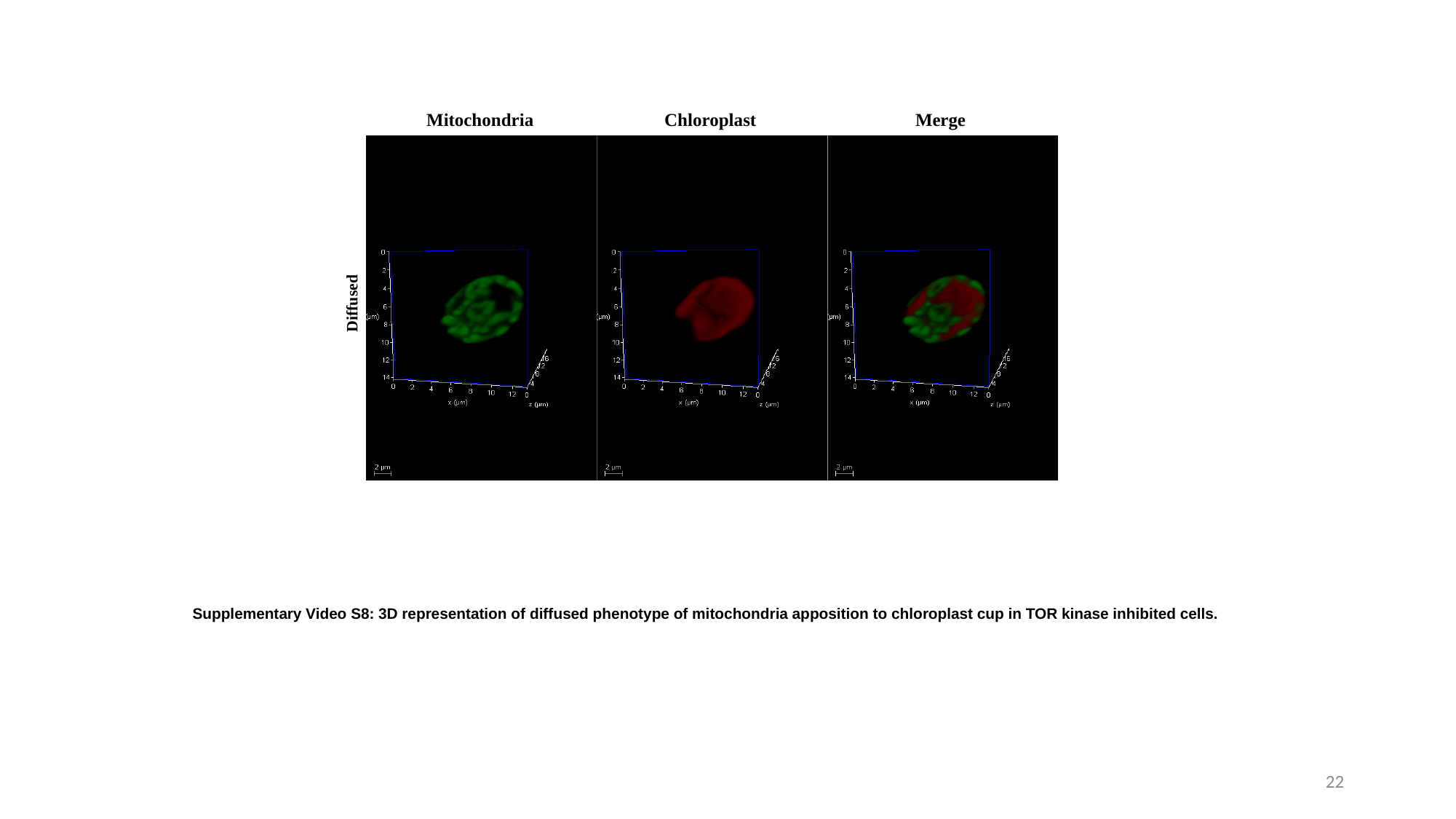

Mitochondria
Chloroplast
Merge
Diffused
Supplementary Video S8: 3D representation of diffused phenotype of mitochondria apposition to chloroplast cup in TOR kinase inhibited cells.
22
